# Red light perception by the root is essential for gibberellin-induced primary-root elongation in tomato

**DOI:** 10.1101/2023.08.20.554010

**Authors:** Uria Ramon, Amit Adiri, Hadar Cheriker, Ido Nir, Yogev Burko, David Weiss

## Abstract

- The promoting effect of gibberellin (GA) on primary-root elongation is well-documented in several plant species, yet its influence in others, including tomato (*Solanum lycopersicum*), remains unclear.
- The role of GA in primary-root elongation has been studied in tomato using the GA-deficient mutants *gib-1* and *ga20-oxidase* (*ga20ox1*) and various growth systems, including Dark (D)-root and D-shoot plates.
- GA application to these mutants following germination on vermiculite, promoted primary-root elongation. However, when the roots grew deeper into the dark environment the hormone had no effect. RNA-seq analysis of dark-grown roots, treated with GA, revealed typical transcriptional responses, but the output for cell expansion remained unaffected. When dark-grown roots were illuminated deep in the ground, the hormone promoted their elongation. The results suggest that activation of Phytochrome B (PhyB) in the root, by red light, is essential for GA-induced elongation.
- We propose that GA promotes tomato root elongation after germination, when roots are exposed to low light underground and this contributes to rapid seedling establishment. As roots penetrate deeper into the soil, insensitivity to GA due to the lack of light may be important for sustained root growth under fluctuating water availability, given that water deficiency suppresses GA accumulation.

## Introduction

The plant hormone gibberellin (GA) regulates numerous developmental processes, including organ elongation (Yamaguchi, 2008; Hedden & Sponsel, 2015). Bioactive GAs bind to the GIBBERELLIN-INSENSITIVE DWARF1 (GID1) receptor, leading to the interaction of GID1s with the master growth suppressor, the DELLA protein (Locascio *et al*., 2013). This facilitates the degradation of DELLA by the 26S proteasome, a process mediated by SCF^SLY^. DELLA degradation leads to transcriptional reprogramming and activation of GA responses, including shoot and root elongation (Hauvermale *et al*., 2012).

The role of GA in primary root elongation has been studied mainly in Arabidopsis (*Arabidopsis thaliana*) and its central role was demonstrated (Ubeda-Tomas *et al*., 2008, 2009; Achard *et al*., 2009; Rizza *et al*., 2017). GA regulates both cell division in the root meristem (Shtin *et al*., 2022), and cell elongation in the root elongation zone (Ubeda-Tomas *et al*., 2008). The Arabidopsis GA deficient mutant *ga1-3* has short primary root. Normal elongation can be restored in this mutant by GA application or loss of DELLA activity (Fu & Harberd, 2003). Baker *et al*. (2021) showed that roots are autonomous for GA production and its biosynthesis occurs mainly in the cortex and the endodermis. Using the GA perception sensor nlsGPS1, Rizza *et al*. (2017) showed that active endogenous GA accumulates in the root elongation zone. Ubeda-Tomas *et al*. (2008) showed that inhibition of GA signaling specifically in the endodermis is sufficient to disrupt root elongation, indicating that the endodermis is the key site for GA action in the regulation of root elongation. This conclusion was supported by Shani *et al*., (2013) that found accumulation of bioactive tagged-GAs in the endodermis of the root elongation zone. While root elongation in the GA deficient mutant *ga1-3* was induced by low concentration of GA, this treatment had no effect on shoot growth (Arizumii *et al*., 2008), suggesting that roots are more sensitive to GA than shoot. Moreover, it has been demonstrated that very low concentrations of GA are sufficient to saturate the root elongation response in GA deficient mutant (Tanimoto & Hirano, 2013).

The importance of GA in the regulation of root elongation in some other plant species is less clear, with numerous conflicting reports (Torrey, 1976; Feldman, 1984; Phinney, 1985; Tanimoto, 2005; Tanimoto & Hirano, 2013; Ramon *et al*., 2020). While Whaley and Kephart (1957) show that GA application to maize (*Zea mays*) promotes root elongation, Svensson (1972) reported that the hormone has no effect on maize root elongation. In Medicago (*Medicago truncatula*), GA treatment suppressed, and the GA-biosynthesis inhibitor paclobutrazol (PAC) increased primary root elongation (Fonouni-Farde *et al*., 2019), suggesting that GA inhibits Medicago root elongation. Gou *et al*. (2010) showed in *Populus* trees that reduced GA activity promotes lateral root elongations and has no effect on primary-root length. Butcher & Street (1960) found that elevating GA concentrations in tomato (*Solanum lycopersicum*), progressively promote root elongation, whereas Tognoni *et al*. (1967), showed that application of GA to tomato plants inhibits root elongation in a concentration-dependent manner.

In a previous study we showed in tomato seedlings that PAC treatment slightly inhibited primary root elongation (Ramon *et al*., 2020). Application of low GA concentrations to the PAC-treated seedlings restored root elongation, but higher concentrations, inhibited it, exhibiting a bell-shaped curve of response. In contrast, in the shoots, GA strongly promoted elongation and the effect was saturated at very high concentrations and exhibited a typical saturated-type curve of response. Genetic evidence also supports a promotive role for GA in tomato primary root elongation. This is exemplified by the shorter primary root of the strong GA-deficient mutant *gib-1* (Barlow *et al*., 1991), which lacks the activity of ent-copalyl diphosphate synthase (CPS, Koornneef *et al*., 1990; Bensen & Zeevaart, 1990). Similarly, the roots of the extreme dwarf mutant *gid1^TRI^*, which lack the activity of all GA receptors, are shorter (Illouz-Eliaz *et al*., 2019; Ramon *et al*., 2020). Although these studies show a clear effect for the inhibition of GA synthesis/activity on tomato root elongation, the effect of these mutations on root growth was much weaker than their effect on stem elongation (Ramon et al., 2020).

Most studies on GA and root elongation were performed with light-grown seedlings on agar plates (Arizumii *et al*., 2008; Ubeda-Tomas *et al*., 2008; Shani *et al*., 2013; Rizza *et al*., 2017). Roots however, normally grow in the soil, in the dark and escape from light by negative phototropism (Zhang *et al*., 2013). Using the Dark-root (D-root) system, Silva-Navas *et al*. (2015) showed that roots respond differently to many stimuli under light or dark conditions. Here we have studied how GA affects tomato root elongation underground, and why its effect is much weaker on root elongation than on stem elongation. We demonstrated that activation of Phytochrome B (PhyB) in the root tissue, by red light, is essential for the promoting effect of GA on primary root elongation. The results suggest that penetration of light to the upper layers of the soil drives this GA response, but later, when the roots grow dipper into the dark environment, GA has no more effect on root elongation.

## Materials and Methods

### Plant materials, seed germination, growth conditions and hormone treatments

Tomato cv. M82 (sp/sp), *gib-1* (Koornneef *et al*., 1990; Bensen & Zeevaart, 1990) and *ga 20-oxidase* (*ga20ox1*, Shohat *et al*., 2021) in M82 background and *phyB1phyB2* double mutant (Weller *et al*., 2000) in Money Maker (MM) background were used in this study. Plants/seedlings were grown in vermiculite in a growth room set to a photoperiod of 12 h/12 h, day/night, light intensity of 150μmol m^−2^ s^−1^ and 25°C. In other experiments, plants were grown in a glasshouse under natural day-length conditions, a light intensity of 700– 1000 µmol m^−2^ s^−1^ and 18–30°C. The tomato seeds were sterilized with ethanol 70% for 2 minutes, and then with sodium hypochlorite 3% for 10 minutes. The seeds washed thoroughly by sterilized double distillated water (DDW). For germination of the strong GA deficient mutant *gib-1*, embryos were rescued as described by Illouz-Eliaz et al., (2019) and for germination of *ga 20-oxidase1* (*ga20ox1*, Shohat *et al*., 2021) mutant seeds were scarified.

### Root elongation assay in Petri dishes (agar plates)

The tomato seedlings were grown in Petri dishes filled with 4.4 g/L Murashige & Skoog (MS) medium (Sigma-Aldrich), 8 g/L plant agar (Duchefa Biochemie, RV Haarlem, The Netherlands), pH adjusted to 5.7 with NaOH (with or without 10µM GA_3_, 10 mg/L PAC or both). The plates were placed vertically under continuous white light (100μmol m^−2^ s^−1^, provided by cool-white lamps). For dark condition the plates were covered with aluminum foil after germination. In some experiments (when germination was not uniform), the location of the root tip was marked on the plate following germination, and 5-7 days later, the length from the mark to the root tip was measured. For length measurements the plates were photographed (Canon PowerShot SX60 HS, Melville, NY, USA) and hypocotyl and primary-root length was measured using ImageJ software (http://rsb.info.nih.gov/ij).

### D-root and D-shoot growth system

To grow seedlings on agar plates (see above) and keep the root in the dark and the shoot in the light, we have used a modified D-root growth system (Silva-Navas et al., 2015; Supporting information Fig. S1). We used black plastic comb that was fitted into the plates separating the shoot and the root system and covered the lower part (root part) of the plate with aluminum foil. To keep the shoot in the dark while exposing the root to the light we used Dark-shoot (D-shoot) growth system. We used the same system as for D-root, but covered the upper part (shoot part) of the plate with aluminum foil. The plates were placed vertically under white light (100μmol m^−2^ s^−1^, provided by cool-white lamps).

### Light treatments

Seedlings were grown on D-shoot plates in a growth chamber (Percival, Perry, Iowa, USA), at 25°C, under the specified constant light conditions: blue light (19μmol m^−2^ s^−1^), red light (8μmol m^−2^ s^−1^), and far-red light (17μmol m^−2^ s^−1^). Light for all experiments was provided by light-emitting diodes (LED30-HL1). The wavelengths and light intensity were measured by LI-180 Spectrometer (LI-COR Biosciences, Lincoln, NE, USA) (Supporting information methods S2).

### Hormonal treatments

GA_3_, and/or PAC were applied by spraying and drenching when grown on vermiculite and incorporated into the growth medium, when grown on agar plates.

### Molecular Cloning/Constructs and Plant Transformation

To generate *p35S:GID1a* construct, *GID1a* (Illouz-Eliaz *et al*., 2019) cDNA was inserted into the pART7 vector downstream to the 35S promoter into the KpnI and XbaI sites. The construct was subcloned into the pART27 binary vector by NotI site and introduced into *Agrobacterium tumefaciens* strain GV3101 by electroporation. The construct was transferred into M82 cotyledons using transformation and regeneration methods described before (McCormick, 1991). Kanamycin-resistant T0 plants were grown and at least seven independent transgenic lines were selected and self-pollinated to generate homozygous transgenic lines.

### Library Constructions and RNAseq analysis

Total RNA was extracted from root tips and leaves using a RNeasy Plant Mini Kit (Qiagen). RNA-seq libraries were prepared at the Crown Genomics institute of the Nancy and Stephen Grand Israel National Center for Personalized Medicine, Weizmann Institute of Science, Israel. Libraries were prepared using the INCPM-mRNA-seq protocol. Briefly, the polyA fraction (mRNA) was purified from 500ng of total input RNA followed by fragmentation and the generation of double-stranded cDNA. After Agencourt Ampure XP beads cleanup (Beckman Coulter), end repair, a base addition, adapter ligation, and PCR amplification steps were performed. Libraries were quantified by Qubit (Thermo Fisher Scientific, Waltham, MA 15 USA) and TapeStation (Agilent, Santa Clara, CA, USA). Sequencing was done on a NextSeq instrument (Illumina, San Diego, CA, USA) using a HO 75 cycles kit, allocating approximately 18 million reads per sample (single read sequencing).

### Sequence data analysis

Stretches of Poly-A/T and Illumina adapters were trimmed from the reads using cutadapt (Martin *et al*., 2011); resulting reads shorter than 30bp were discarded. Reads were mapped to the Solanum lycopersicum reference genome; release SL3.00 using STAR (Dobin *et al.,* 2013) (with End To End option and out Filter Mismatch Nover Lmax set to 0.04). Annotation file was downloaded from SolGenomics, ITAG release 3.2. Expression levels for each gene were quantified using htseq-count (Anders *et al*., 2014), using the gtf file above (using union mode). Differentially expression analysis was performed using DESeq2 (Love *et al*., 2014) with the betaPrior, cooksCutoff and independent Filtering parameters set to False. Raw P values were adjusted for multiple testing using the procedure of Benjamini and Hochberg (Bejamini & Hochberg, 1995). The RNA-Seq data discussed in this publication have been deposited in NCBI’s Gene Expression Omnibus (Edgar *et al*., 2002):

https://dataview.ncbi.nlm.nih.gov/object/PRJNA1002623?reviewer=m2cghdn5fcve5npf4 qpqd8es63.

### RNA Extraction, cDNA Synthesis and RT-qPCR Analysis

Total RNA was extracted with a RNeasy Plant Mini Kit (Qiagen, Hilden, Germany). For synthesis of cDNA, SuperScript II reverse transcriptase (18064014; Invitrogen, Waltham, MA, USA) and 3 mg of total RNA were used according to the manufacturer’s instructions. RT-qPCR analysis was performed using an Absolute Blue qPCR SYBR Green ROX Mix kit (AB-4162/B; Thermo Fisher Scientific, Waltham, MA, USA). Reactions were performed using a Rotor-Gene 6000 Cycler (Corbett Research, Sydney, Australia). A standard curve was obtained using dilutions of the cDNA sample. The expression was quantified using the Rotor-Gene’s software (Corbett Research). Three independent technical repeats were performed for each sample. Relative expression was calculated by dividing the expression level of the examined gene by that of *SlACTIN*. The target-gene-to-*ACTIN* ratio was then averaged.

### Confocal Imaging and cell length measurement

Root tips were stained with Propidium iodide (PI) solution (0.01μg ml^–1^). Laser beam of 552 nm were used for excitation of PI. The samples were placed on a glass slide with a drop of water and sealed with coverslip. Fluorescence images were acquired using Leica SP8 confocal microscope (Boston, MA, USA). The cells lengths were measured by ImageJ software (http://rsb.info.nih.gov/ij).

### Statistical analysis

All assays were conducted with three or more biological replicates and analyzed using JMP software (SAS Institute, Cary, NC, USA). Means comparison was conducted using analysis of variance (ANOVA) with post-hoc Tukey–Kramer honest significant difference (HSD) test (for multiple comparisons) and Student’s t-test (for one comparison). For assays without normal distribution Mann-Whitney -Wilcoxon test was conducted. The experiments were performed three times.

### Accession numbers

Sequence data from this article can be found in the Sol Genomics Network (https://solgenomics.net/) under the following accession numbers: ACTIN, Solyc11g005330; GA20ox1, Solyc03g006880; GA20ox3, Solyc11g072310; GID1a, Solyc01g098390; GID1b1, Solyc09g074270

## Results

To study the effect of GA on root elongation in tomato, we first treated WT M82 seedlings, grown in vermiculite with 10μM GA_3_. A week later, we measured primary root length but found no significant effect for the hormone (Supporting information Fig. S3). This imply that the effect of endogenous GA on primary root elongation is saturated in WT plants, similar to Arabidopsis (Tanimoto & Hirano, 2013). Thus, we investigated the impact of GA on primary root elongation, in the strong GA-deficient mutant *gib-1*, known for its heigh sensitivity to exogenous GA (Bensen & Zeevaart, 1990; Koornneef *et al*., 1990). This mutant exhibited very low percentage of germination even after seed scarification, and therefore, we had to rescue its embryos. *gib-1* and WT (M82) rescued embryos were planted on vermiculite, in pots, and after the seedlings produced four expanded leaves, the length of the primary roots was measured. Similar to *gid1^TRI^* (Ramon *et al*., 2020), primary roots of *gib-1* were shorter than those of WT (Fig. 1a). Microscopic analysis revealed that epidermal cells of *gib-1* primary roots are shorter than those of WT roots (Supporting information Fig. S4). We then examined the effect of GA application to *gib-1* on primary-root elongation. Two weeks-old *gib-1* seedlings, grown in vermiculite, were treated (spraying and drenching) twice a week for one month with 10μM GA_3_ and then the length of leaves, stems and primary roots was measured. While the hormone strongly promoted shoot growth, it had no effect on primary root elongation (Fig. 1b,c). We further examined the effect of GA on primary-root elongation in a weaker GA-deficient mutant *ga20ox1* (Shohat *et al*., 2021). This CRISPR-derived mutant germinates following seed scarification and has shorter primary root than the WT M82 (Supporting information Fig. S5a). Scarified *ga20ox1* seeds were planted on vermiculite and 14 days post-germination the seedlings were treated with 10μM GA_3_ and a week later the length of the primary roots was measured. GA had a very small promoting effect on *ga20ox1* primary root elongation (Supporting information Fig. S5b).

**Fig. 1.**
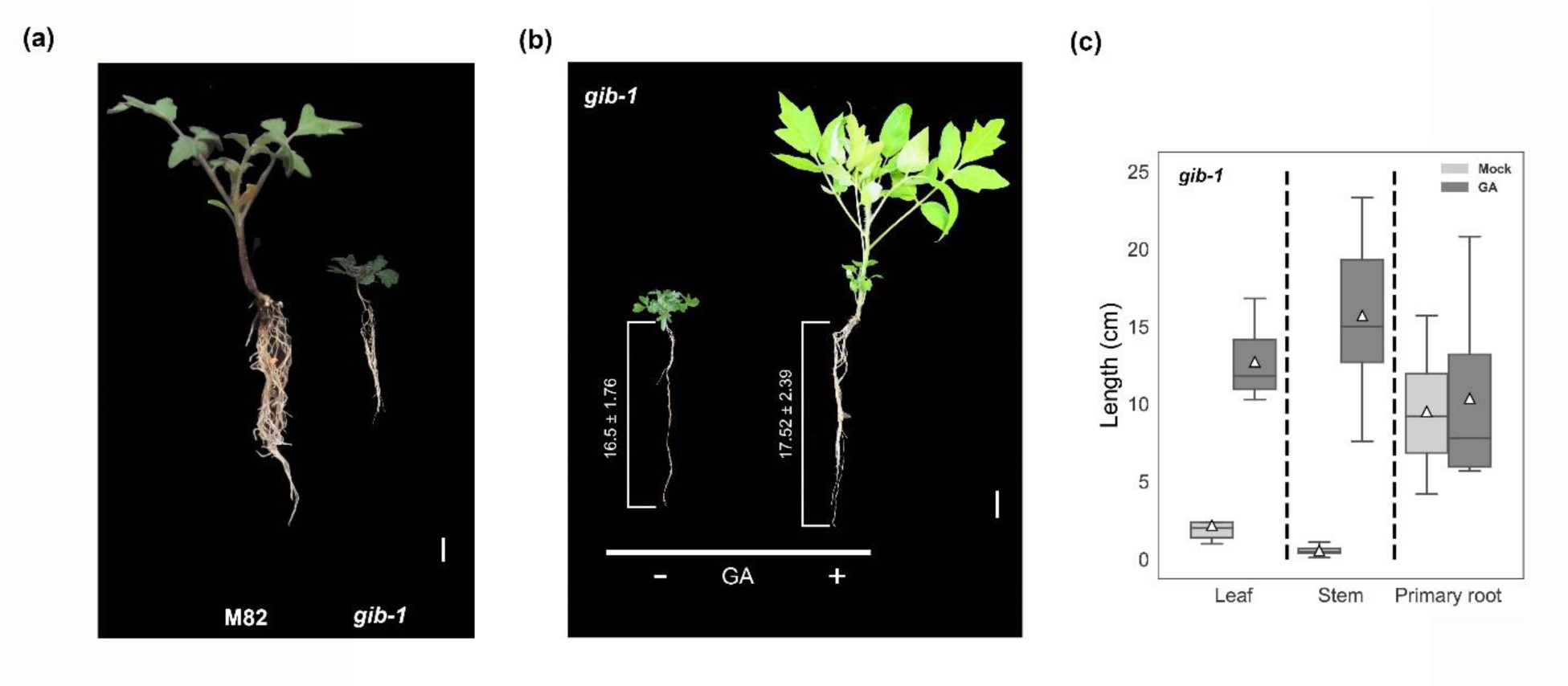
GA application to *gib-1* had no effect on primary-root elongation. (a). Representative WT M82 and *gib-1* plants at the same physiological age (three leaves). (b and c) Two weeks-old *gib-1* seedlings, grown in vermiculite, were treated twice a week for one month with 10μM GA_3_ by dranching. (b) Representative GA-treated and non-treated *gib-1* plants. The average length of the primary roots is presented (n=10 ±SE). (c) Leaf (third leaf from the top), stem and primary root length of GA treated or non-treated *gib-1* plants. Data are graphically presented as whisker and box plots (n=10 ±SE). White triangles show the average values (n=8 ±SE). Bars in a = 1 cm in b = 2.5 cm.

As demonstrated above, the impact of exogenous GA on primary root elongation was minimal, if any, for both *gib-1* and *ga20ox1*. We hypothesized that GA is not active in the roots, deeper in the growth medium. We therefore, examined general and specific GA transcriptional responses in these roots. We grew WT and *gib-1* for two weeks on vermiculite, and then treated *gib-1* plants with 10μM GA_3_ by drenching and spraying and collected shoots and primary roots 24 h later for RNAseq analysis. A previous study in Arabidopsis showed that after a short period from the GA treatment (1–3 h), growth-related genes were only slightly affected by the hormone, but after 24 h, these genes exhibited clear upregulation (Park et al., 2017). We first analyzed the expression of all canonical components involved in GA signaling in WT shoots and roots. The dominant GA receptor *GID1a* (Illouz-Eliaz *et al*., 2019) exhibited similar expression in shoots and roots (Fig. 2a). The other two receptors, *GID1b1* and *GID1b2*, exhibited much higher expression in the roots than in the shoot. The F-box *SLEEPY1* (*SLY1*) showed similar expression in shoot and roots and the tomato *DELLA*, *PROCERA,* exhibited higher expression in the shoot (Fig. 2a). These results suggest that all GA signaling components are expressed in the root. In a previous study (Ramon et al., 2020) we speculated that in tomato, root elongation is less responsive to GA than the shoot due to lower activity of GID1a, whose activity is responsible for the strong stem-elongation response to GA (Illouz-Eliaz *et al*., 2019). The results of the RNAseq do not support this hypothesis. Moreover, overexpression of *GID1a* in M82 roots slightly inhibited their elongation and eliminated completely the effect of GA (Supporting information Fig. S6), probably due to super-optimal GA activity in the root (Ramon *et al*., 2020).

**Fig 2.**
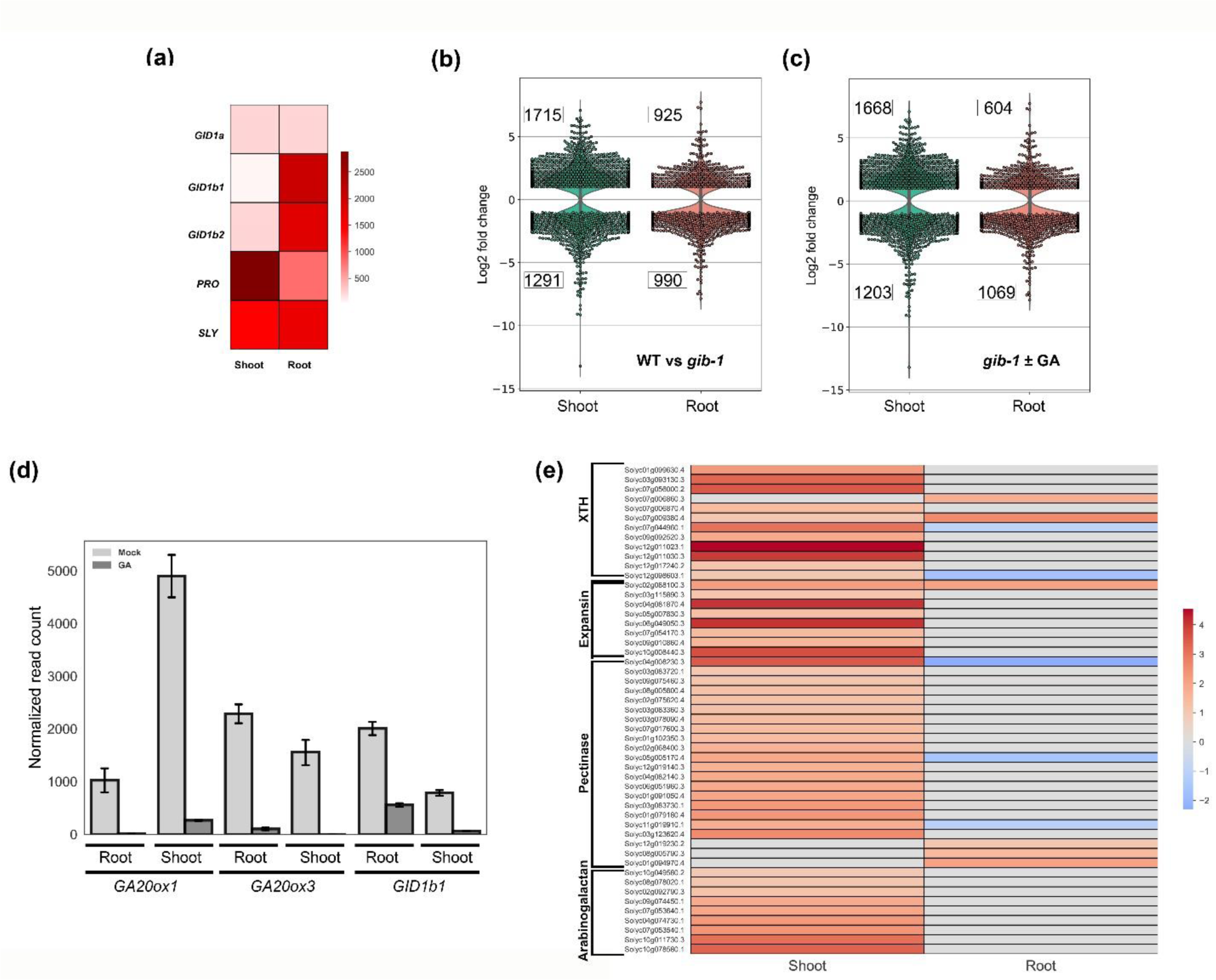
Transcriptional response to GA in tomato root and shoot. M82 and *gib-1* seedlings were grown for two weeks on vermiculite, and then *gib-1* plants were treated by drenching and spraying with 10μM GA_3_ and 24 h later, shoots and primary root were taken (three biological replications) for RNAseq analysis. (a) Heatmap showing normalized read counts of GA-signaling genes in WT M82 shoots and roots. Values are the average of three biological repeats. Coloring of the genes is according to the color bar (normalized read counts).(b and c) Violine plots of differentially expressed genes (DEG) between (b) WT M82 and *gib-1* shoot and root. (c) shoots and roots of *gib-1* treated or not treated with GA. (d) Normalized read counts of *GA20ox1*, *GA20ox3* and *GID1b1* in *gib-1* roots and shoots treated or not treated with GA. Values are the average of three biological repeats ±SE. (e) Heatmap of GA regulated cell expansion genes (Park et al., 2017) in *gib-1* shoots and roots. Coloring of the genes is according to the color bar (Log2 fold change). XTH-xyloglucan endotransglucosylase.

We next compared non-treated *gib-1* and WT shoot and roots. In shoots we found ca. 3000 differentially expressed genes (DEGs) between the two genotypes and in roots approximately 1900 DEG (Fig. 2b, Supporting information Fig. S7a, Supporting information Dataset S1). When *gib-1* was treated with GA, both the shoots and the roots exhibited clear transcriptional responses to the treatment; approximately 2900 DEG were identified in the shoot and 1700 DEGs in the roots (Fig. 2c, Supporting information Fig. S7b, Supporting information Dataset S1). Moreover we found the typical negative feedback response to GA in the roots; the expression of *GA20ox1, GA20ox3* and *GID1b1* was strongly inhibited in *gib-1* shoots and roots by the GA treatment (Fig. 2d). The activation of the feedback and the general transcription response suggest active GA signaling and responses in the roots. We then analyzed the expression of cell-expansion genes in *gib-1* shoots and roots following GA treatment (Park et al., 2017). While many of these genes, including *EXPANSINs*, *XYLOGLUCAN ENDO-TRANSGLYCOSYLASEs* (*XTHs*), *PECTINASEs* and *ARABINOGALACTANs* were upregulated following the GA treatment in the shoots, in the roots they were hardly affected (Fig. 2e). Collectively, these results suggest that although GA signaling and most response pathways are active in dark-grown roots, the GA-output pathway leading to cell elongation remained inactive.

Numerous studies demonstrating a significant impact of GA on primary root elongation were conducted on agar plates under light conditions (Ubeda-Tomas *et al*., 2008, 2009; Achard *et al*., 2009; Rizza *et al*., 2017; Qin *et al*., 2022). We therefore tested if GA can promote elongation of *gib-1* primary root in the light. Rescued *gib-1* embryos were grown on agar plates with or without 10μM GA_3_ and the plates were placed vertically in the light or under dark conditions for five days. GA induced hypocotyl elongation in both light and dark (Fig. 3a and b), but promoted primary root elongation only in the light (Fig. 3a and c). We also examined microscopically the length of the primary-root epidermal cell above the elongation zone and found that GA increased cell length in the light but had no effect on cell elongation in the dark (Supporting information Fig. S8). To study if GA is active in these dark grown roots, we analyzed the feedback response to GA in *gib-1* roots grown in agar plates under light or dark conditions with or without GA. The expression of *GA20ox1, GA20ox3* and *GID1b1* genes was strongly inhibited by GA in both light- and dark-grown roots (Fig. 3d), supporting our hypothesis that although GA is active in dark-grown roots, the output toward root elongation is blocked. We further examined the effect of light vs. dark on GA-induced primary root elongation in *ga20ox1* seedlings and found similar results: GA promoted primary root elongation only in the light (Supporting information Fig. S9). GA had very mild promoting effect on M82 primary root elongation under light conditions, implying that the GA level in WT roots is close to saturation for elongation (Supporting information Fig. S10).

**Fig. 3.**
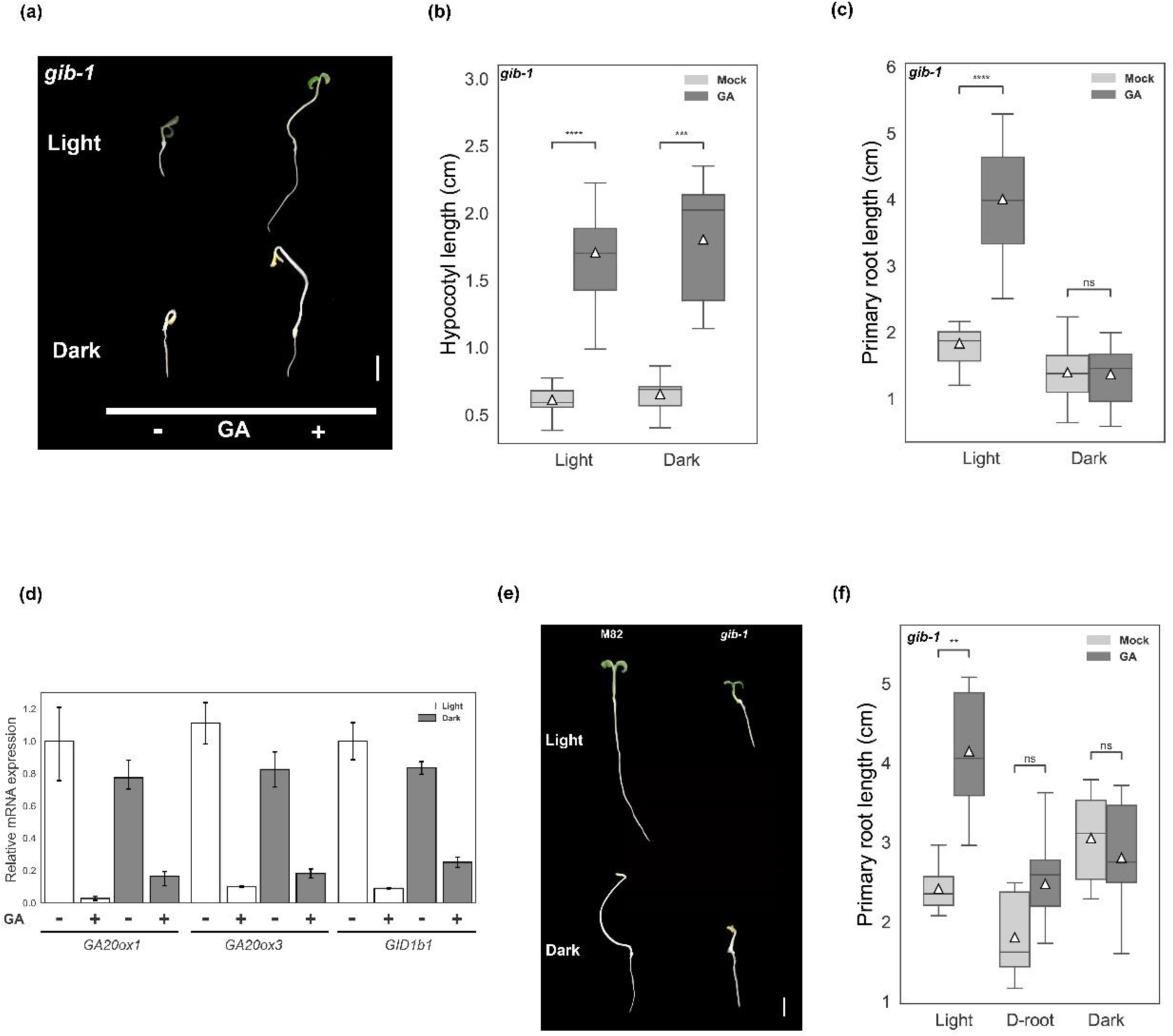
Light is required for the promoting effect of GA on primary-root elongation. (a, b and c) Seedlings (*gib-1*) were grown in agar plates with or without 10μM GA_3_ (incorporated in the growth medium) and the plates were placed vertically in the light or under dark conditions and after 5 days hypocotyl and primary-root length was measured. (a) Representative seedlings (bars = 1 cm). (b) Hypocotyl length. (c) Primary root length. Data in b and c are graphically presented as whisker and box plots (n=16) ± SE. White triangles show the average values. (d) RT-qPCR analyses of *GA20ox1*, *GA20ox3* and *GID1b1* expression in *gib-1* roots grown as described above. Values (relative expression) are means of 4 biological replicates ±SE. The value of mock treated light-grown roots for each gene was set to 1. (e) Representative M82 and *gib-1* seedlings grown in agar plates for 5 days in light or dark (bar = 1 cm). (f) Seedlings (*gib-1*) were grown in agar plates or D-root plates with or without 10μM GA_3_ and the plates were placed vertically in the light (agar plates and D-root plates) or in dark (agar plates only). After 5 days the length of the primary roots was measured. Values are mean of 15 seedlings ±SE. Stars in b, c, d and f represent significant differences between respective treatments by Student’s t test (*P < 0.05; **P<0.01; ***P<0.001). ns- not significant.

If roots are insensitive to GA in the dark, then under dark conditions the length of M82 roots should be similar to those of *gib-1*. We grew M82 and *gib-1* on agar plates under light or dark conditions. M82 hypocotyls were much longer than those of *gib-1* under light and dark conditions. However, M82 primary roots were longer than those of *gib-1* only under light conditions and under dark conditions their length was similar (Fig. 3e, Supporting information Fig. S11). To test whether direct light to the root is essential for GA-induced elongation, we utilized the D-root agar plate system (Silva-Navas et al., 2015; Supporting information Fig. S1). This system allows seedlings to grow on agar plates with their shoots exposed to light while the roots develop in the dark, resembling the conditions experienced by soil-grown seedlings. While GA promoted *gib-1* primary root elongation in the light, it had a very mild effect on root elongation in the D-root system (Fig. 3f). It should be noted that the D-root system does not block completely the light to the root, but just limiting it (Silva-Navas *et al*., 2015).

We hypothesized that GA induces root elongation right after germination, when light penetrates the upper layer of the soil (Tester & Morris, 1987; Mandoli *et al*., 1990), but later, deeper in the soil, in a complete darkness, the roots cannot respond to GA. To test this hypothesis, we applied 10μM GA_3_ immediately after planting the rescued *gib-1* embryos on the vermiculite, or 10 days later, when the roots were deeper (approximately 2 cm long) in the vermiculite. While GA_3_ application immediately after sowing promoted primary root elongation, at the later stage the hormone had no effect (Fig. 4a). Moreover, a single application of GA (10μM GA_3_) right after germination completely restored *gib-1* root elongation and their primary-root length, two weeks later, was similar to that of the WT (Supporting information Fig. S12). To eliminate the possibility that root sensitivity to GA is restricted to early growth stages and diminishes with time, we examined the root elongation of *gib-1* seedlings grown on agar plates in the presence or absence of GA_3_, or initially without GA followed by the addition of GA_3_ only when primary roots reached approximately 2.5 cm in length, all under light conditions. GA promoted elongation when applied at early or later stages (Fig. 4b), ruling out the possibility that GA can induce elongation only at the very early stages of primary-root development, and further support the hypothesis that perception of light by the root promotes GA-induced elongation.

**Fig. 4.**
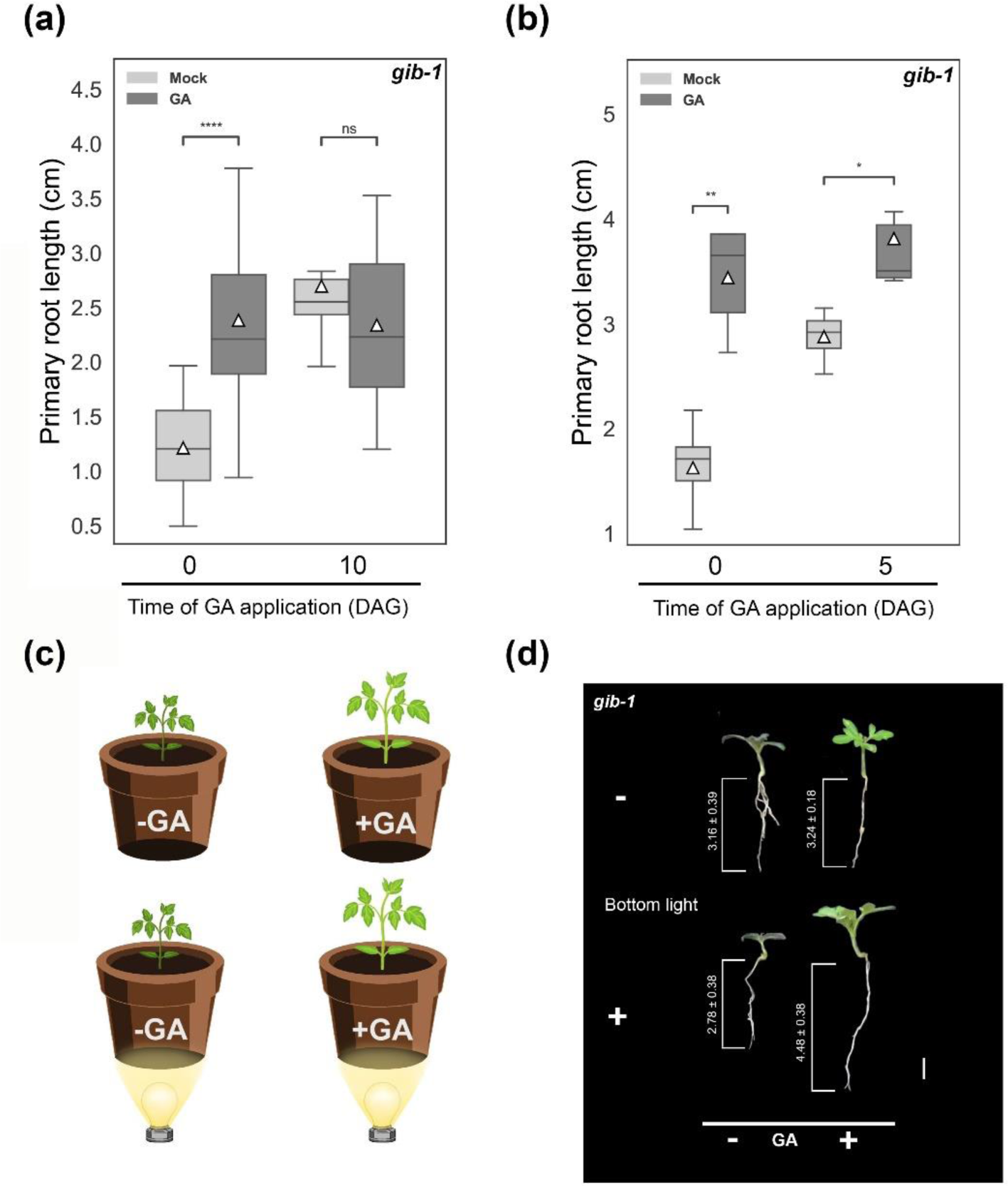
The promoting effect of GA on primary root elongation right after germination required light and was induced deeper in the ground by direct illumination. (a) Rescued *gib-1* embryos were placed on vermiculite and treated by drenching with 10μM GA_3_ immediately, or 10 days later. One week after the GA treatment (for each treatment), the length of the primary roots was measured. Data are graphically presented as whisker and box plots (n= 16 ± SE). (b) Rescued *gib-1* embryos were planted in agar plates without or with 10μM GA_3_ in the growth medium or first without GA and only when primary roots were approximately 2.5 cm long, 10μM GA_3_ solution was added for 30 min and then removed. The seedlings were grown in the light for 6 days and then the length of the primary roots was measured. Data are graphically presented as whisker and box plots (n= 12 ± SE). Stars represent significant differences between respective treatments or lines by Student’s t test (*P < 0.05; **P<0.01; ***P<0.001; ****P<0.0001). ns- not significant. DAG- days after germination. White triangles in a and b show the average values. (c) The bottom lighting experimental setup. (d) *gib-1* embryos were planted on vermiculite in bottomless pots that were placed (or not) on LED lighting strips and two weeks later, when roots were approximately 2.5 cm long, the light was turned on and half of the plants were treated by drenching with 10μM GA_3_. A week later, the length of the primary roots was measured for all treatments. Representative plants with the average length of the primary roots (n=16 ±SE) are presented in the picture. Bar= 1 cm.

To exclude the possibility that the lack of effect for GA on root elongation deeper in the ground is caused by the mechanical resistant of the soil (Colebrook *et al*., 2014) and not by lack of light, we examined if the effect of GA on root elongation could be induced by light deeper in the soil. For this purpose, we removed the bottoms of the pots, and the pots containing vermiculite were placed over LED lighting strips, to expose root to light from the bottom (Fig. 4c and Supporting information Fig. S13). *gib-1* embryos were planted on the vermiculite and two weeks later, when roots were approximately 2.5-3 cm long (approximately 1.5-2 cm from the bottom), the light was turned on and GA_3_ was applied by drenching. A week later, we measured the length of the primary roots of all treatments. GA application to LED-lighted *gib-1* roots promoted their elongation, but without bottom lighting it had no effect (Fig 4d and Supporting information Fig. S14), supporting our hypothesis that direct light to the root is essential for GA-induced primary root elongation.

If GA-induced primary root elongation in the upper layer of the soil, we anticipated that the activation of GA-induced elongation will be responsive to low light intensity. We examined the effect of light intensity on GA-induced hypocotyl and primary-root elongation. Rescued *gib-1* embryos were placed on agar plates with or without GA_3_ in dark or under 0.1, 1, 10 or 100µmol m^-2^ s^-1^ of white light. The effect of GA_3_ on hypocotyls elongation was found in both dark and light and was not affected by light intensity (Fig 5a and b and Supporting information Fig. S15a). On the other hand, GA had no effect on primary-root elongation in the dark, but a clear effect was found already at the lowest light intensity (0.1 µmol m^-2^ s^-1^) and the strongest effect was found under 100 µmol m^-2^ s^-1^ (Fig 5a and b and Supporting information Fig. S15b).

**Fig. 5.**
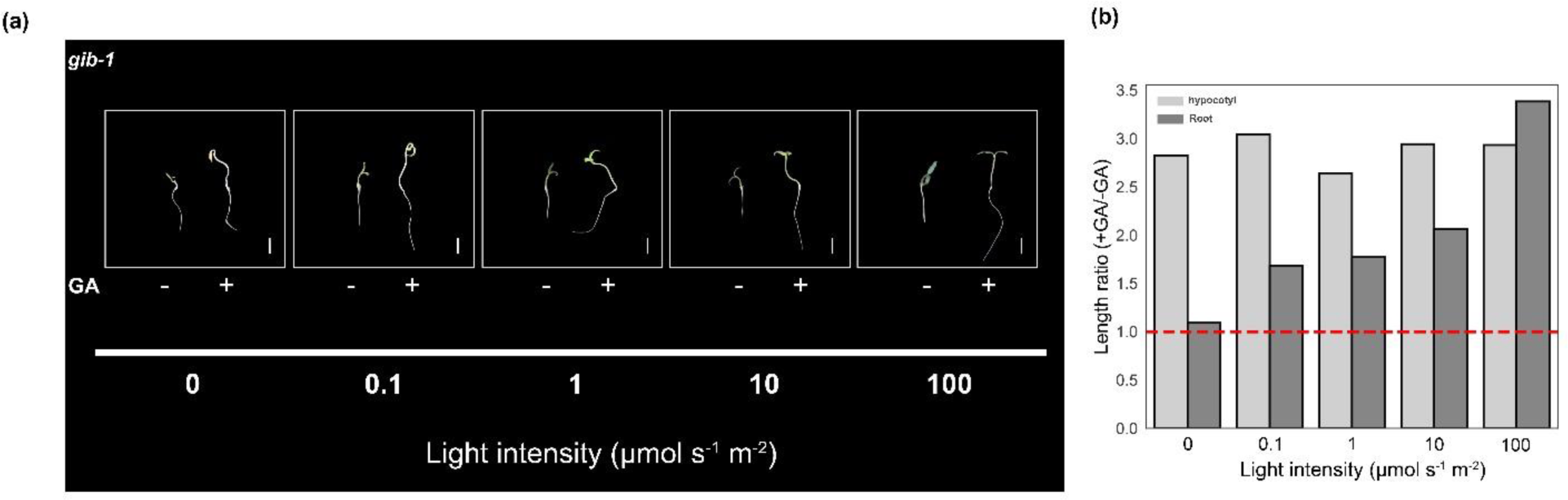
Low light intensity is sufficient for GA to promote primary root elongation. Rescued *gib-1* embryos were placed on agar plates with or without GA_3_ in the growth medium in the dark or under 0.1, 1, 10 or 100 µmol m^-2^ s^-1^ of white light. After 10 days hypocotyl and primary root length was measured. (a) Representative seedling. Bars = 1 cm. (b) The length ratio between GA treated and non-treated seedlings (hypocotyl and primary roots). Values are mean of 9 seedlings.

Analysis of our RNAseq data revealed that the photoreceptors *PhyA*, *PhyB1*, *PhyB2*, *Cryptochrome 1* (*CRY1*), *CRY1b*, *CRY2*, *Phototropin 1* (*Phot1*) and *Phot2* are all expressed in tomato roots (Fig. 6a). To study the specific light spectrum essential for GA-induced root elongation and the specific photoreceptor/s involved, we employed a novel growth system that allows direct light exposure to the root while maintaining the shoot in darkness (D-shoot, see Supporting information Fig. S1). GA stimulated primary root elongation when the roots were exposed to light (D-shoot), but not when the roots grew in the dark in D-root plates (Supporting information Fig. S16). Subsequently, we utilized the D-shoot system to explore the active light spectrum. Rescued *gib-1* embryos and scarified *ga20ox1* seeds were planted in D-shoot agar plates with or without GA_3_ under blue, red, or far-red light (see methods for light spectrum details). GA promoted primary root elongation in both mutants only under red light, but not under blue or far-red light (Fig. 6b and c). Considering that the effect of continuous red light is mediated by PhyB (Quail, 2002), we tested if mutations in the two *PhyB* genes (*phyB1 phyB2* double mutant, Weller et al., 2000) would inhibit the effect of GA on primary-root elongation under light conditions. To this end, *phyB1 phyB2* and its WT background MM seeds were placed first in agar plates under white light and right after germination the young seedlings were transferred to D-shoot agar plates, containing 10 mg/L PAC to reduce the endogenous level of GA, or PAC with 10μM GA_3_ and the plates were placed under continuous red or white light. GA promoted primary root elongation in MM under red and white light but had no effect under the same growth conditions on *phyB1 phyB2* primary root elongation (Fig 6d), suggesting that light-induced GA elongation response is mediated by PhyB activity in the root.

**Fig. 6.**
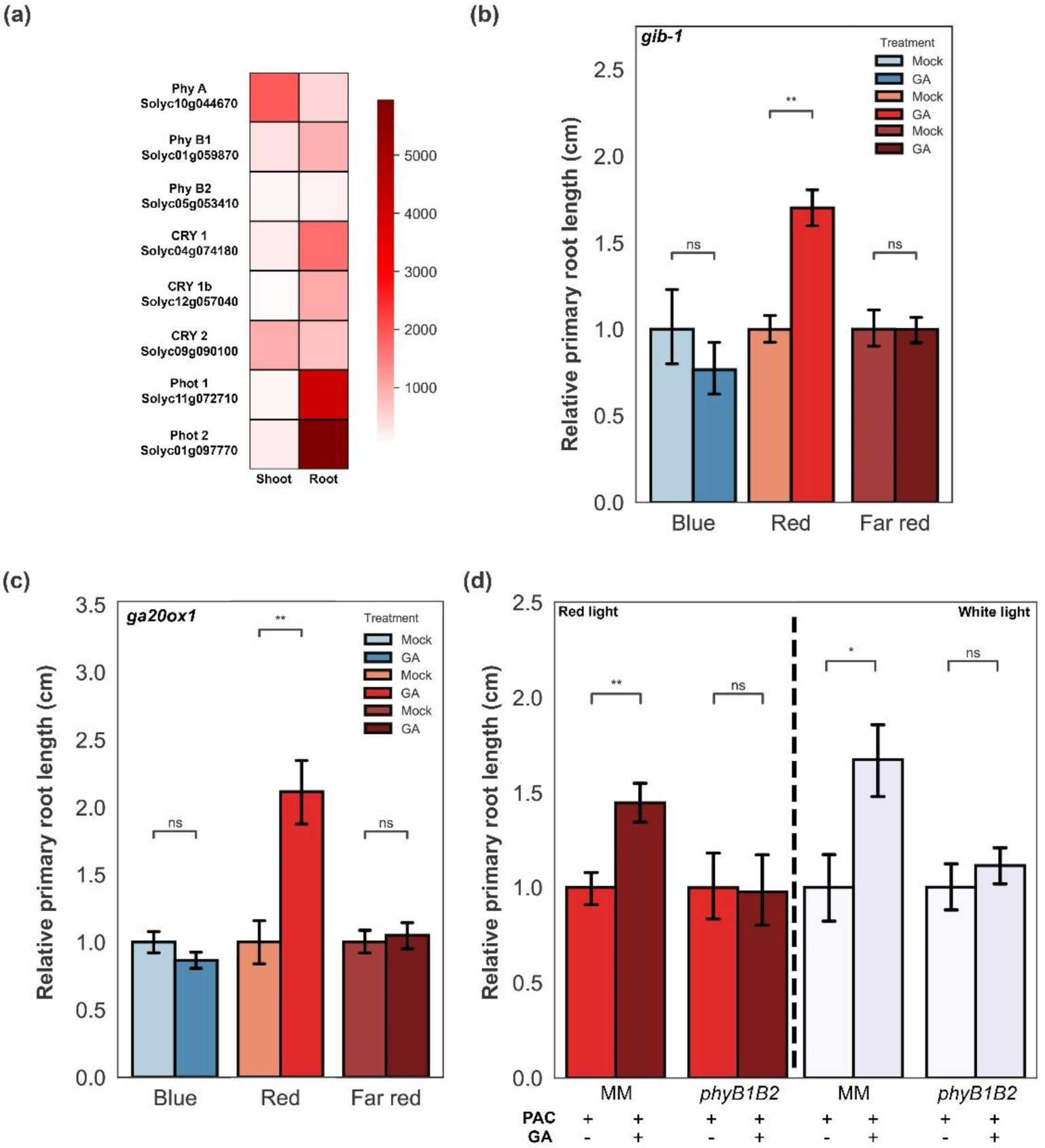
Activation of PhyB in the root by red light is required for GA to promote primary root elongation. (a) Normalized reads count of the different photoreceptors (with their accession numbers) in tomato M82 shoot and root. The data were taken from the RNAseq analysis (see above). (b and c) *gib-1* (rescued embryos) and *ga20ox1* scarified seeds were planted on D-shoot plates with or without 10μM GA_3_ in the growth medium under blue, red or far-red light (for light spectrum see Supporting information Fig. S2) for 5 days and then primary-root length was measured. (d) WT MM and *phyB1phyB2* double mutant seedlings were grown in D-shoot plates with 10mg/l PAC or PAC and 10μM GA_3_ in the growth medium under red or white light for 5 days and then primary-root length was measured. The mean values of the PAC treatment in b, c and d were set to 1 and those of the PAC+GA treatment are the ratio between PAC+GA/PAC mean. The values in b and c are mean of 16 seedlings and in d, 25 seedlings ±SE. Stars represent significant differences between respective treatments by Student’s t test (P<0.01). ns- not significant.

## Discussion

Rapid elongation of the primary root is crucial for seedling establishment, as it anchors the seedling in the soil after germination and facilitates immediate water and nutrient uptake. Several factors regulate this rapid, directional elongation, including environmental factors, such as light and gravity that guide growth downward into the soil (negative phototropism and gravitropism), as well as the interplay of growth hormones. Auxin plays a central role in root growth, but other hormones, including brassinosteroids and GA are also known to promote root elongation (Garay-Arroyo *et al*., 2012). The role of auxin has been studied intensively and is conserved in all tested plant species, but the role of GA in different species is less clear. Numerous studies have demonstrated a clear promoting role for GA in primary root elongation, but others have indicated no effect or even an inhibitory effect (Tognoni *et al*. 1967; Tanimoto & Hirano 2013; Fonouni-Farde *et al*., 2019; Castro-Camba *et al*., 2022). The disparities among these reports may arise from the application of GA to WT plants with already saturated GA levels in the root, in some studies, or from the application of the hormone to GA-deficient mutants, in other studies. Additionally, the variation could be attributed to the use of low GA concentrations, promoting elongation in certain studies, and the application of super-optimal GA concentrations inhibiting root elongation in others (Inada & Shimmen, 2000; Ramon *et al*., 2021). Our study suggests that the experimental setup may also contribute to these discrepancies, whether the plants are grown on plates under light conditions or in soil with roots growing in the dark. Silva-Navas *et al*. (2015) demonstrated that roots behave differently under light and dark conditions. Our results further indicate that in tomato, the promoting effect of GA on primary root elongation is influenced by light. Consequently, the hormone’s effect would differ when examined on agar plates under light conditions or in the soil in the dark.

The interplay between light and GA was demonstrated in various developmental processes, including germination, hypocotyl and stems elongation and root development. According to these studies, light affects GA metabolism, transport and responses (Arana *et al*., 2014; Matsuo *et al*., 2019; Lyu *et al*., 2021; Li *et al*., 2020; van Gelderen *et al*., 2023). In soybean (*Glycine max*), blue light, via CRY, suppresses stem elongation by the induction of *GA 2-oxidase* (*GA2ox*) transcription, leading to GA deactivation (Lyu *et al*., 2021). In Arabidopsis activated CRY (by blue light) interacts with DELLA, inhibits its polyubiquitination and degradation and therefore, suppresses GA responses, including hypocotyl elongation (Xu *et al*., 2021). Low red/far-red ratio, mediated by PhyB, affects GA accumulation and shoot elongation in *Pinus tabuliformis* seedlings (Li *et al*., 2020). It also affects GA responsiveness in Arabidopsis seeds (Arana *et al*., 2014). Elevated levels of far-red light directed towards Arabidopsis shoots stimulate the accumulation of GA and its subsequent translocation to the roots, where it acts to inhibit lateral root growth (van Gelderen *et al*., 2023). Our findings reveal that exposure of primary root to continuous red light, but not to blue or far-red light, promotes GA-induced primary root elongation, indicating a role for PhyB (Quail, 2002). Indeed, GA had no effect on primary root elongation in *phyB1 phyB2* double mutant, under red or white light.

The dependency of GA-induced primary root elongation on red light can explain, at least partially, why the effect of GA deficiency on tomato root elongation is much weaker than the effect on stem elongation, as previously demonstrated (Ramon et al., 2020). Since light penetrates only to the upper layer of the ground, the effect of the hormone is restricted to this area and just for the initial growth phase of the root. Later, in the dark, the process of root elongation becomes insensitive to GA. In the stem on the other hand, GA continuously promotes elongation.

The mechanism by which red light, via PhyB, affects GA-induced root elongation responses in tomato is yet unknow. In dark-grown roots, all GA signaling components were expressed and GA induced or suppressed the expression of numerous genes and pathways. Nevertheless, the hormone had no effect on cell expansion genes (Fig. 2). The activation of the feedback response by GA (down regulation of *GA20ox1*, *GA20ox3* and *GID1b1*) in dark grown roots, is an indication for DELLA degradation by GA (Fukazawa *et al*., 2017). This suggests that light promotes GA-induced root-elongation response downstream to DELLA. DELLA, a key regulator of GA responses, modulates downstream pathways by interacting with numerous transcription factors (TF) and either inhibiting or promoting their activity (Locascio *et al*., 2013). The degradation of DELLA by GA in dark-grown roots releases these TFs and probably triggers transcriptional activation of numerous pathways independently of light. These may include the regulation of root architecture (Castro-Camba *et al*., 2022), secondary xylem development, lignification (Wang et al., 2017) and endodermal suberin formation (Binenbaum *et al*., 2023), but not cell elongation. Several DELLA-interacting TFs are involved in GA regulation of cell elongation (de Lucas *et al*., 2008; Feng *et al*., 2008; Gallego-Bartolome *et al*., 2012; Chaiwanon & Wang, 2015). Based on the presented results, it is reasonable to suggest that in tomato roots the activity of a DELLA-interacting, elongation-related TF, and/or its downstream pathway are regulated by light

Roots normally grow in the dark and escape from light by negative phototropism (Zhang *et al*., 2013). However, in most cases, shortly after germination, in the upper layer of the soil, the young roots are exposed to low light intensity (Tester & Morris, 1987; Mandoli *et al*., 1990). We suggest that in tomato, at this early stage the light signal which is mediated by PhyB interacts with the GA response pathway to accelerate root elongation. Rapid elongation at this early stage of root development contributes to seedling establishment. At later stages however, when the root elongation zone is deeper in the soil, in the dark, GA have no more effect on root elongation. It is possible that the insensitivity of root elongation to GA deeper in the soil, contributes to root growth under changing environment. Under moderate drought conditions, bioactive GA levels in tomato decrease due to the inhibition of GA biosynthesis and increased GA deactivation (Shohat *et al*., 2021). The reduced GA levels suppresses shoot growth to reduce water loss. At the same time, under the same dry conditions, root elongation is maintained (xerotromism), enabling roots to grow deeper and find new sources of water (Sharp *et al*., 2004; Miao *et al*., 2021). If root elongation was sensitive to changes in GA levels, this differential growth response between shoot and root, under moderate drought conditions, would not be possible.

## Acknowledgements

This research was supported by the Israel Science Foundation (617/20) to DW. We thank Dr. Natanella Illouz-Eliaz for her valuable suggestions. We would like to thank Matar Azriel for the illustration in Figure 4. We also thank Ziva Amsellem for technical assistance.

## Author Contributions

UR, IN, YB and DW designed the research; UR, HC, and AA performed experiments; UR, AA and DW analyzed data; UR, IN, YB and DW wrote the manuscript.

## Data Availability

All data can be found in the manuscript and in the supporting information.

## References

Achard P, Gusti A, Cheminant S, Alioua M, Dhondt S, Coppens F, Beemster GTS, Genschik P. 2009. Gibberellin signaling controls cell proliferation rate in Arabidopsis. Current Biology 19: 1188–1193

Anders S, Pyl PT, Huber W. 2015. HTSeq—a Python framework to work with high-throughput sequencing data. bioinformatics, 31: 166–169.

Arana MV, Sánchez-Lamas M, Strasser B, Ibarra SE, Cerdán PD, Botto JF, Sánchez RA. 2014. Functional diversity of phytochrome family in the control of light and gibberellin-mediated germination in Arabidopsis. Plant Cell & Environment 37: 2014–2023.

Ariizumi T, Murase K, Sun T, Steber CM. 2008. Proteolysis-Independent Downregulation of DELLA repression in Arabidopsis by the gibberellin receptor GIBBERELLIN INSENSITIVE DWARF1. The Plant Cell 20: 2447–2459.

Barker R, Fernandez Garcia MN, Powers SJ, Vaughan S, Bennett MJ, Phillips AL, Thomas SG, Hedden P. 2021. Mapping sites of gibberellin biosynthesis in the Arabidopsis roottip. New Phytologist 229: 1521–1534.

Barlow PW, Brain P, Parker JS. 1991. Cellular growth in roots of a gibberellin-deficient mutant of tomato (*Lycopersicon escolentum* Mill.) and its wild-type. Journal of Expiramental Botany 42: 339–351.

Bensen RJ, Zeevaart JAD. 1990. Comparison of *ent*-kaurene synthetase A and B activities in cell-free extracts from young tomato fruits of wild-type and *gib-1*, *gib-2*, and *gib-3* tomato plants. Journal of Plant Growth Regulation 9: 237–242.

Binenbaum J, Wulff N, Camut L, Kiradjiev K, Anfang M, Tal I, Vasuki H, Zhang Y, Sakvarelidze-Achard L, Davière JM, et al. 2023. Gibberellin and abscisic acid transporters facilitate endodermal suberin formation in Arabidopsis. Nature Plants 9: 785– 802

Benjamini Y, Hochberg Y. 1995. Controlling the false discovery rate: a practical and powerful approach to multiple testing. Journal of the Royal Statistical Society 57: 289–300.

Butcher DN, Street HE. 1960. Effects of kinetin on the growth of excised tomato roots. Physiologia Plantarum 13: 46–55.

Castro-Camba R, Sánchez C, Vidal N, Vielba JM. 2022. Plant development and crop yield: the role of gibberellins. Plants 11(19), 2650

Chaiwanon J, Wang Z-Y. 2015. Spatiotemporal brassinosteroid signaling and antagonism with auxin pattern stem cell dynamics in Arabidopsis roots. Current Biology 25: 1031–1042

Colebrook EH, Thomas SG, Phillips AL, Hedden P. 2014. The role of gibberellin signalling in plant responses to abiotic stress. Journal of Experimental Biology 217: 67–75.

de Lucas M, Daviére JM, Rodríguez-Falcón M, Pontin M, Iglesias-Pedraz JM, Lorrain S, Fankhauser C, Blázquez MA, Titarenko E, Prat S. 2008. A molecular framework for light and gibberellin control of cell elongation. Nature 451: 480–484.

Dobin A, Davis CA, Schlesinger F, Drenkow J, Zaleski C, Jha S, Gingeras TR. 2013. STAR: ultrafast universal RNA-seq aligner. Bioinformatics 29: 15–21.

Edgar R, Domrechev M, Lash AE. 2002. Gene Expression Omnibus: NCBI gene expression and hybridization array data repository. Nucleic Acids Research 30: 207–210.

Feldman LJ. 1984. Regulation of root development. Annual review of plant physiology 35: 223–242.

Feng S, Martinez C, Gusmaroli G, Wang Y, Zhou J, Wang F, et al. 2008. Coordinated regulation of *Arabidopsis thaliana* development by light and gibberellins. Nature 451: 475–479.

Fonouni-Farde C, Miassod A, Laffont C, Morin H, Bendahmane A, Diet A, Frugier F. 2019. Gibberellins negatively regulate the development of *Medicago truncatula* root system. Scientific Reports 9: 1–9.

Fu X, Harberd NP. 2003. Auxin promotes Arabidopsis root growth by modulating gibberellin response. Nature 421: 740–743.

Gallego-Bartolome J, Minguet, EG, Grau-Enguix F, Abbas M, Locascio A, Thomas SG, Alabadi D, Blazquez MA. 2012. Molecular mechanism for the interaction between gibberellin and brassinosteroid signaling pathways in Arabidopsis. Proceedings of the National Academy of Sciences 109: 13446–13451.

Fukazawa J, Mori M, Watanabe S, Miyamoto C, Ito T, Takahashi Y. 2017. DELLA-GAF1 complex is a main component in gibberellin feedback regulation of GA20 oxidase 2. Plant Physiology 175: 1395–1406

Garay-Arroyo A, De La Paz Sánchez M, Garcıá-Ponce B, Azpeitia E, Álvarez-Buylla ER. 2012. Hormone symphony during root growth and development. Developmental Dynamics 241:1867–1885

van Gelderen K, van der Velde K, Kang C-k, Hollander J, Petropoulos O, Akyüz T, Pierik R. 2023. Gibberellin transport affects (lateral) root growth through HY5 during Far-Red light enrichment. bioRxiv doi: 10.1101/2023.04.21.537844

Gou J, Strauss SH, Tsai CJ, Fang K, Chen Y, Jiang X, Busov VB. 2010. Gibberellins regulate lateral root formation in populous through interactions with auxin and other hormones. The Plant Cell 22: 623–639

Hauvermale AL, Ariizumi T, Steber CM. 2012. Gibberellin signaling: a theme and variations on DELLA repression. Plant Physiology 160: 83–92.

Hedden P, Sponsel V. 2015. A Century of Gibberellin Research. Journal of Plant Growth Regulation 34: 740–760

Illouz-Eliaz N, Ramon U, Shohat H, Blum S, Livne S, Mendelson D, Weiss D. 2019. Multiple gibberellin receptors contribute to phenotypic stability under changing environments. The Plant Cell 31: 1506–1519.

Inada S, Shimmen T. 2000. Regulation of elongation growth by gibberellin in root segments of *Lemna minor*. Plant and Cell Physiology 41: 932–939.

Koornneef M, Bosma TDG, Hanhart CJ, Veen H Van Der, Zeevaart JAD. 1990. The isolation and characterization of gibberellin-deficient mutants in tomato. Theoretical and Applied Genetics 80: 852–857.

Li W, Liu SW, Ma JJ, Liu HM, Han FX, Li Y, Niu SH. 2020. Gibberellin signaling is required for far-red light-induced shoot elongation in *Pinus tabuliformis* seedlings. Plant Physiology 182: 658–668.

Locascio A, Blázquez MA, Alabadí D. 2013. Genomic analysis of DELLA protein activity. Plant & Cell Physiology 54: 1229–1237.

Love MI, Huber W, Anders S. 2014. Moderated estimation of fold change and dispersion for RNA-seq data with DESeq2. Genome Biology 15:1–21.

Lyu X, Cheng Q, Qin C, Li Y, Xu X, Ji R, Mu R, Li H, Zhao T, Liu J, et al. 2021. GmCRY1s modulate gibberellin metabolism to regulate soybean shade avoidance in response to reduced blue light. Molecular Plant 14: 298–314.

Mandoli DF, Ford GA, Waldron LJ, Nemson JA, Briggs WR. 1990. Some spectral properties of several soil types: implications for photomorphogenesis. Plant Cell & Environment 13: 287–294.

Martin M. 2011. Cutadapt removes adapter sequences from high-throughput sequencing reads. EMBnet Journal 17: 1–10.

Matsuo S, Nanya K, Imanishi S, Honda I, Goto E. 2019. Effects of blue and red lights on gibberellin metabolism in tomato seed. The Horticulture Journal 88: 76–82

McCormick S. 1991. Transformation of tomato with *Agrobacterium tumefaciens*. Plant Tissue Culture Manual B6: 1–9.

Miao R, Yuan W, Wang Y, Garcia-Maquilon I, Dang X, Li Y, Zhang J, Zhu Y, Rodriguez PL, Xu W. 2021. Low ABA concentration promotes root growth and hydrotropism through relief of ABA INSENSITIVE 1-mediated inhibition of plasma membrane H+ATPase 2. Science Advances 7: eabd4113

Park J, Oh D-H, Dassanayake M, Nguyen KT, Ogas J, Choi G, Sun T-p. 2017 Gibberellin signaling requires chromatin remodeler PICKLE to promote vegetative growth and phase transitions. Plant Physiology 173: 1463–1474.

Phinney BO. 1985. Gibberellin A_1_ dwarfism and shoot elongation in higher plants. Biologia Plantarium 27: 172–179.

Qin H, Pandey BK, Li Y, Huang G, Wang J, Quan R, Zhou J, Zhou Y, Miao Y, Zhang, et al. 2022. Orchestration of ethylene and gibberellin signals determines primary root elongation in rice. The Plant Cell 34:1273–1288.

Quail PH. 2002. Phytochrome photosensory signalling networks. Nature Reviews Molecular Cell Biology 3: 85–93.

Ramon U, Weiss D, Illouz-Eliaz N. 2020. Underground gibberellin activity: differential gibberellin response in tomato shoots and roots. New Phytologist 229: 1196–200.

Rizza A, Walia A, Lanquar V, Frommer WB, Jones AM. 2017. In vivo gibberellin gradients visualized in rapidly elongating tissues. Nature Plants 3: 803–813.

Svensson SB. 1972. A comparative study of the changes in root growth, induced by coumarin, auxin, ethylene, kinetin and gibberellic acid. Physiologia Plantarum 26: 115– 135.

Shani E, Weinstain R, Zhang Y, Castillejo C, Kaiserli E, Chory J, Tsien RY, Estelle M. 2013. Gibberellins accumulate in the elongating endodermal cells of Arabidopsis root. Proceedings of the National Academy of Sciences of the United States of America 110: 4834–4839.

Sharp RE, Poroyko V, Hejlek LG, Spollen WG, Springer GK, Bohnert HJ, Nguyen HT. 2004. Root growth maintenance during water defcits: physiology to functional genomics. Journal of Experimental Botany 55: 2343–2351.

Shohat H, Ceriker H, Vasuki H, Illouz-Eliaz N, Blum S, Amsellem Z, Tarkowska D, Aharoni A, Eshed Y, Weiss D. 2021. Inhibition of gibberellin accumulation by water deficiency promotes fast and long-term ‘drought avoidance’ responses in tomato. New Phytologist 232:1985–98.

Shtin M, Dello Ioio R, Del Bianco M. 2022. It’s time for a change: the role of gibberellin in root meristem development. Frontiers in Plant Science 13:882517.

Silva-Navas J, Moreno-Risueno MA, Manzano C, Pallero-Baena M, Navarro-Neila S, Téllez-Robledo B, Garcia-Mina JM, Baigorri R, Gallego FJ, del Pozo JC. 2015. D-Root: a system for cultivating plants with the roots in darkness or under different light conditions. The Plant Journal 84: 244–255.

Stafen CF, Kleine-Vehn J, Maraschin FS. 2022. Signaling events for photomorphogenic root development. Trends in Plant Science 27: 1266–1282.

Tanimoto E. 2005. Regulation of root growth by plant hormones-roles for auxin and gibberellin. Critical Reviews in Plant Sciences 24: 249–265.

Tanimoto E, Hirano K. 2013. Role of gibberellins in root growth. In: Eshel A, Beeckman T. Fourth Edition, Plant Roots: The Hidden Half. CRC Press 13: 1-14

Tester M, Morris C. 1987. The penetration of light through soil. Plant Cell & Environment 10: 281–286

Tognoni F, Halevy AH, Wittwer SH. 1967. Growth of bean and tomato plants as affected by root absorbed growth substances and atmospheric carbon dioxide. Planta 72: 43–52.

Torrey JG. 1976. Root hormones and plant growth. Annual Review of Plant Physiology 27: 435–459.

Ubeda-Tomás S, Swarup R, Coates J, Swarup K, Laplaze L, Beemster GTS, Hedden P, Bhalerao R, Bennett MJ. 2008. Root growth in Arabidopsis requires gibberellin/DELLA signalling in the endodermis. Nature Cell Biology 10: 625–628.

Wang GL, Que F, Xu ZS, Wang F, Xiong AS. 2017. Exogenous gibberellin enhances secondary xylem development and lignification in carrot taproot. Protoplasma 254: 839– 848.

Weller JL, Schreuder ME, Smith H, Kooeneef M, Kendrick RE. 2000. Physiological interactions of phytochromes A, B1 and B2 in the control of development in tomato. The Plant Journal 24: 345-356.

Whaley WG, Kephart J. 1957. Effect of gibberellic acid on growth of maize roots. Science 125: 234.

Xu P, Chen H, Li T, Xu F, Mao Z, Cao X, Miao L, Du S, Hua J, Zhao J, Guo T, Kou S, Wang W, Yang H-Q. 2021. Blue light-dependent interactions of CRY1 with GID1 and DELLA proteins regulate gibberellin signaling and photomorphogenesis in Arabidopsis. The Plant Cell 33: 2375–2394.

Yamaguchi S. 2008. Gibberellin Metabolism and its Regulation. Annual Review of Plant Biology 59: 225–251.

Zhang KX, Xu HH, Yuan TT, Zhang L, Lu YT. 2013. Blue-light-induced PIN3 polarization for root negative phototropic response in Arabidopsis. The Plant Journal 76: 308–321

