## Supporting Information for "Red light perception by the root is essential for gibberellin-induced primary-root elongation in tomato"

***New Phytologist* Supporting Information**

The following Supporting Information is available for this article:

**Fig. S1.** D-root and D-shoot growth systems.

**Fig. S2.** The light spectrum of the different light treatments.

**Fig. S3.** GA treatment has no effect on WT primary root elongation in the soil.

**Fig. S4**. Root epidermal cell length of WT M82 and *gib-1*.

**Fig. S5.** The effect of GA on M82 and *ga20ox1* primary root elongation in the soil.

**Fig. S6.** Effect of *GID1a* overexpression on primary root elongation.

**Fig. S7.** Differentially expressed genes (DEG).

**Fig. S8.** GA promoted primary root cell elongation in the light but not in the dark.

**Fig. S9**. Light is required for GA-induced primary-root elongation in *ga20ox1*..

**Fig. S10.** GA has no effect on primary root elongation in M82 seedlings grown in agar plates.

**Fig. S11.** Primary-root length of WT M82 and *gib-1* grown in agar plates under light or dark conditions.

**Fig. S12.** GA application to *gib-1* immediately after germination on vermiculite restored normal primary root elongation.

**Fig. S13.** The bottom lighting growth system.

**Fig. S14.** Bottom lightening induced the promoting effect of GA on primary root elongation.

**Fig. S15.** GA-induced hypocotyl and primary root elongation under different light intensity.

**Fig. S16.** GA promoted root elongation in D-shoot but not in D-root plates.

**Table S1.** Primers used in this study**.**

**Dataset S1.** Differentially expressed genes (DEG) between M82 and *gib-1* shoots and roots and between *gib-1* shoot and roots, treated or not treated with GA (see separate file).

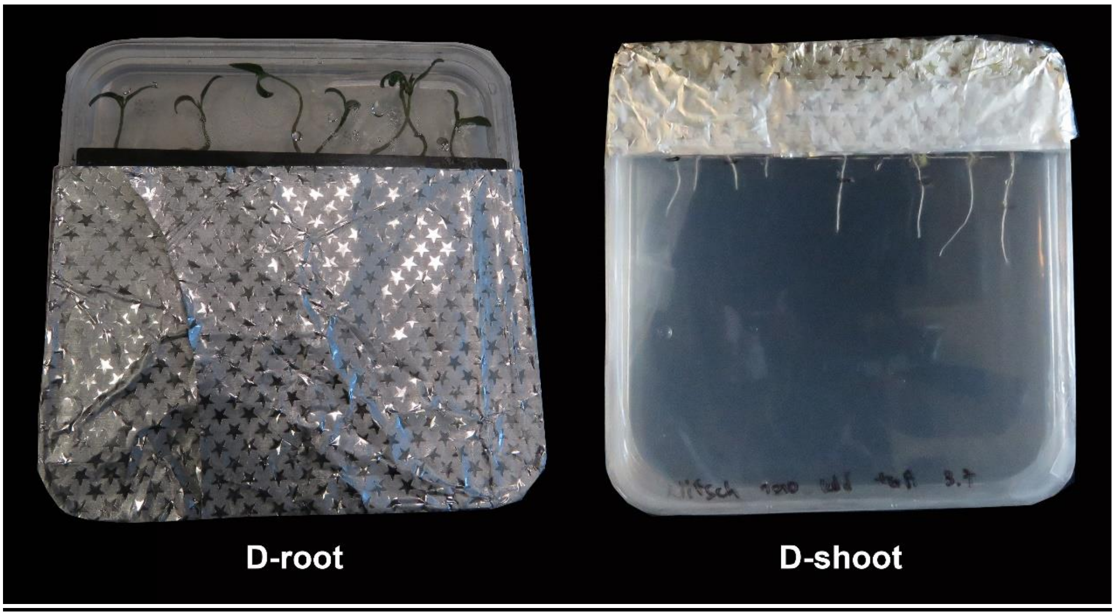


**Fig. S1.** D-root and D-shoot growth systems.

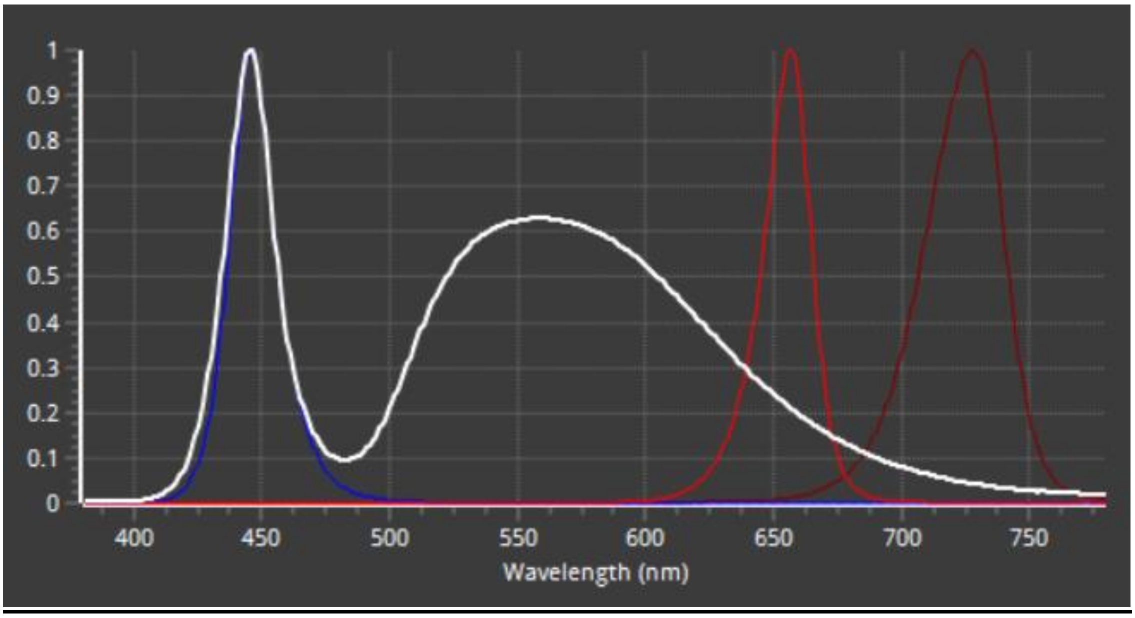


**Fig. S2.** The light spectrum of the different light treatments. White light (white line), blue light (blue line), red light (red line) and far-red light (dark red line).

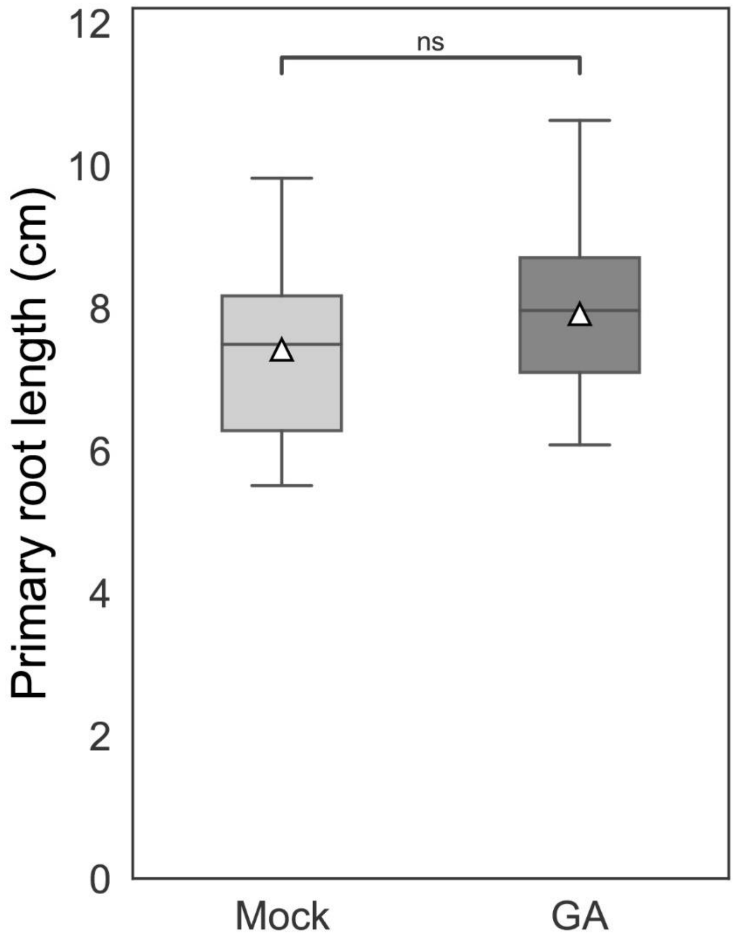


**Fig. S3.** GA treatment has no effect on WT primary root elongation in the soil. M82 seedlings were grown in vermiculite for 10 days and then treated (or not-Mock) with 10μM $GA_3$. A week later primary-root length was measured. White triangles show the average values (n=12 ±SE). ns- not significant by Student's t test.

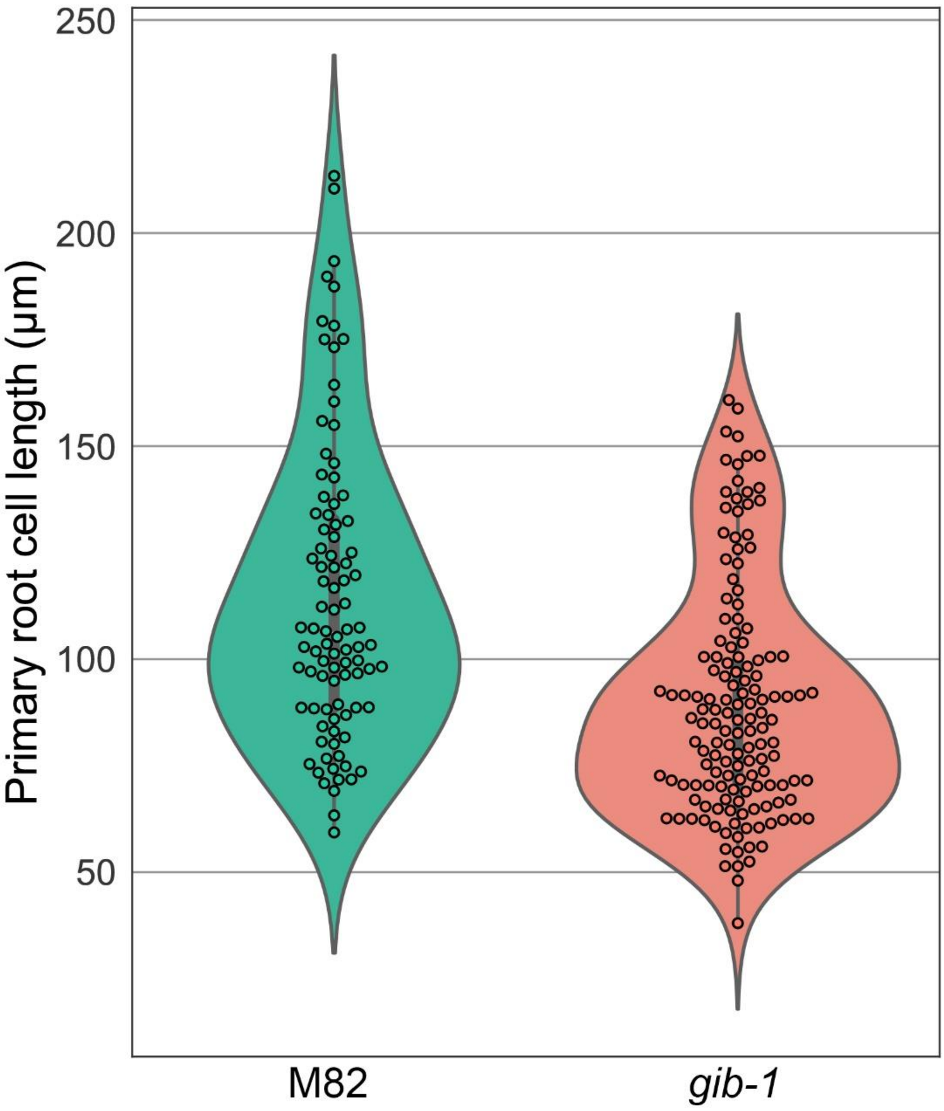


**Fig. S4**. Root epidermal cell length of WT M82 and *gib-1*. Microscopic analysis of epidermal cells above the elongation zone of WT M82 and *gib-1* primary roots.

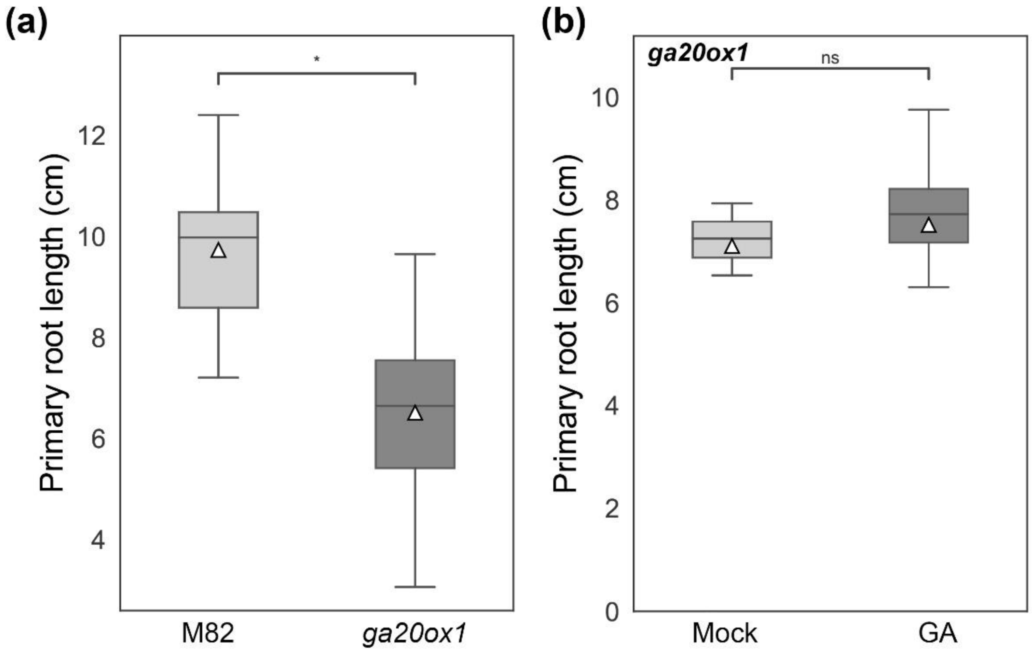


**Fig. S5.** The effect of GA on M82 and *ga20ox1* primary root elongation in the soil. (a) M82 and *ga20ox1* seedlings were grown in vermiculite for 14 days and then primary-root length was measured. (b) *ga20ox1* seedlings were grown for 10 days in the soil and then treated (or not) with 10µM $GA_3$ and 7 days later primary root length was measured. White triangles show the average values (n=7 ±SE). Star represents significant difference by Student's t test (*P < 0.05). ns- not significant.

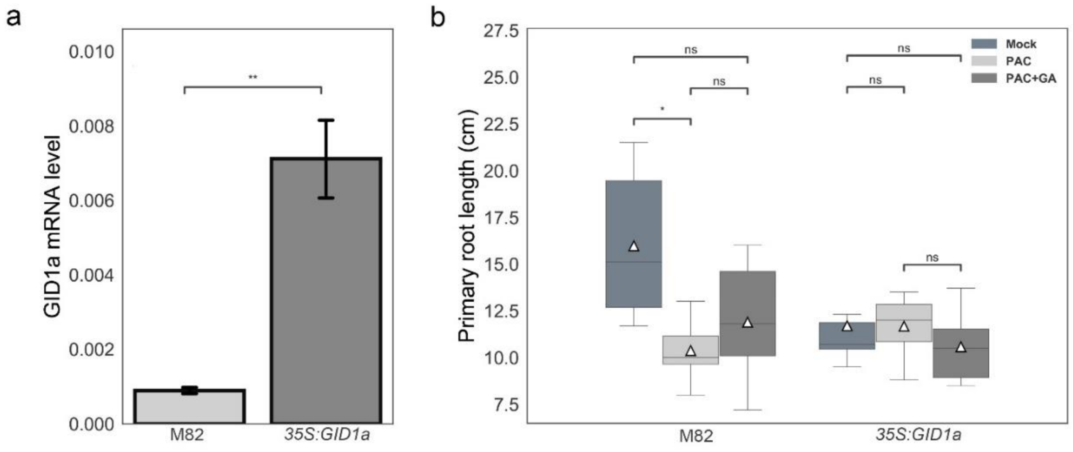


**Fig. S6.** Effect of *GID1a* overexpression on primary root elongation. (a) GID1a expression levels in WT M82 and transgenic *35S:GID1a* roots. Values are mean of three biological replicates ± SE. (b) Primary root length in WT M82 and transgenic plants overexpressing *GID1a* without any treatment or treated with 5mg/l paclobutrazol (PAC) or PAC with 10μM $GA_3$. Data are graphically presented as whisker and box plots (n= 12 ± SE). White triangles show the average values. Stars represent significant differences between respective treatments by Tukey's HSD test (*P<0.05; **P<0.01). ns- not significant.

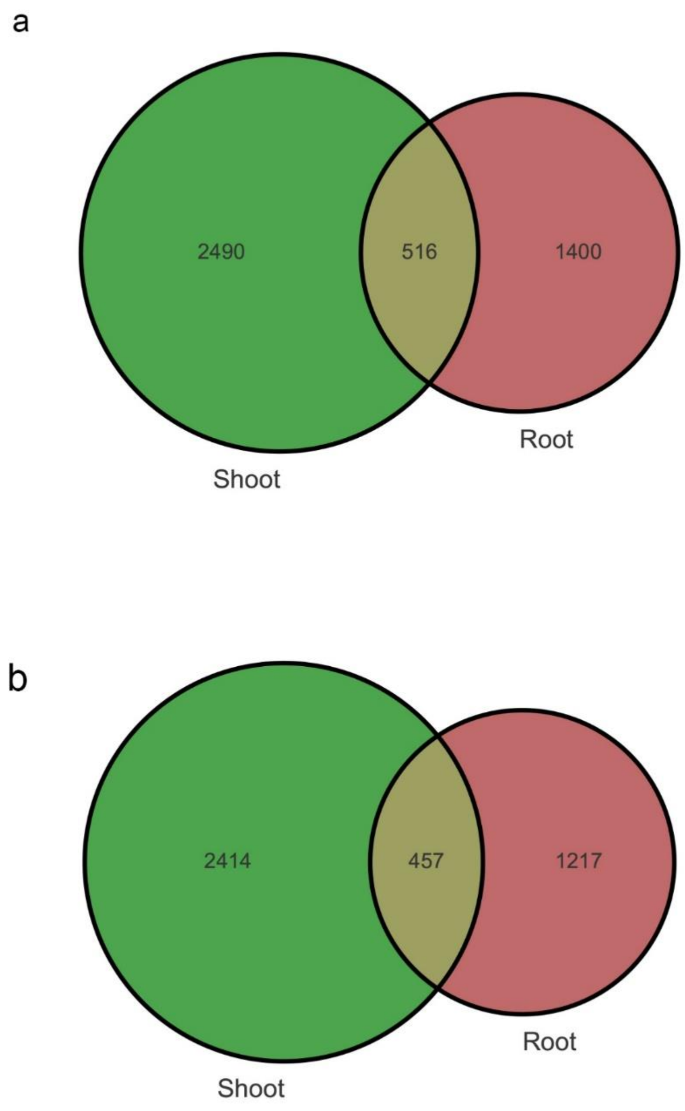


**Fig. S7.** Differentially expressed genes (DEG). (a) Venn diagram represents DEG between WT M82 and *gib-1* shoots and roots. (b) Venn diagram represents DEG between *gib-1* untreated and GA-treated shoots and roots

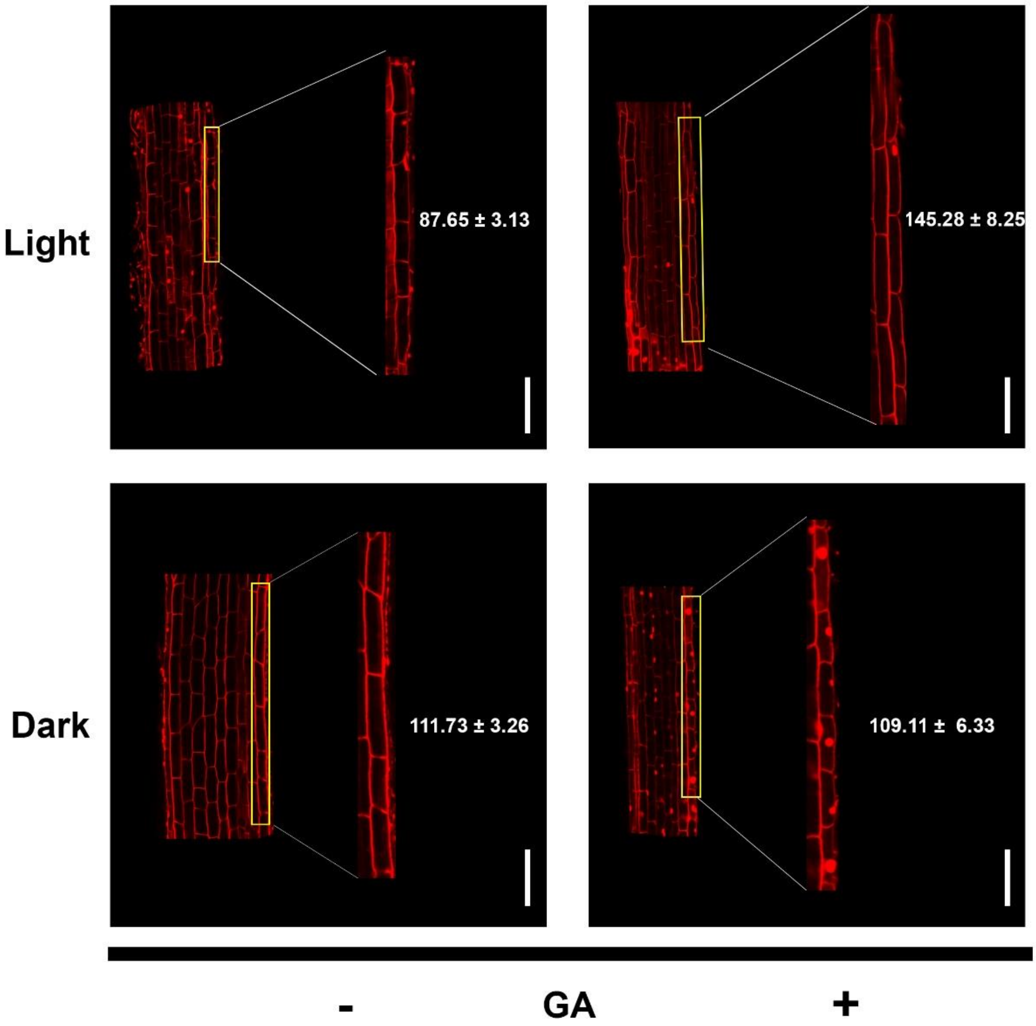


**Fig. S8.** GA promoted primary root cell elongation in the light but not in the dark. *gib-1* seedlings were grown in agar plates with or without 10μM $GA_3$. The plates were placed vertically in the light or in dark conditions for 5 days. The length of the primary root epidermal cell above the elongation zone was examined microscopically. Average cell length is presented (n=30 ± SE). Bar = 100 μm

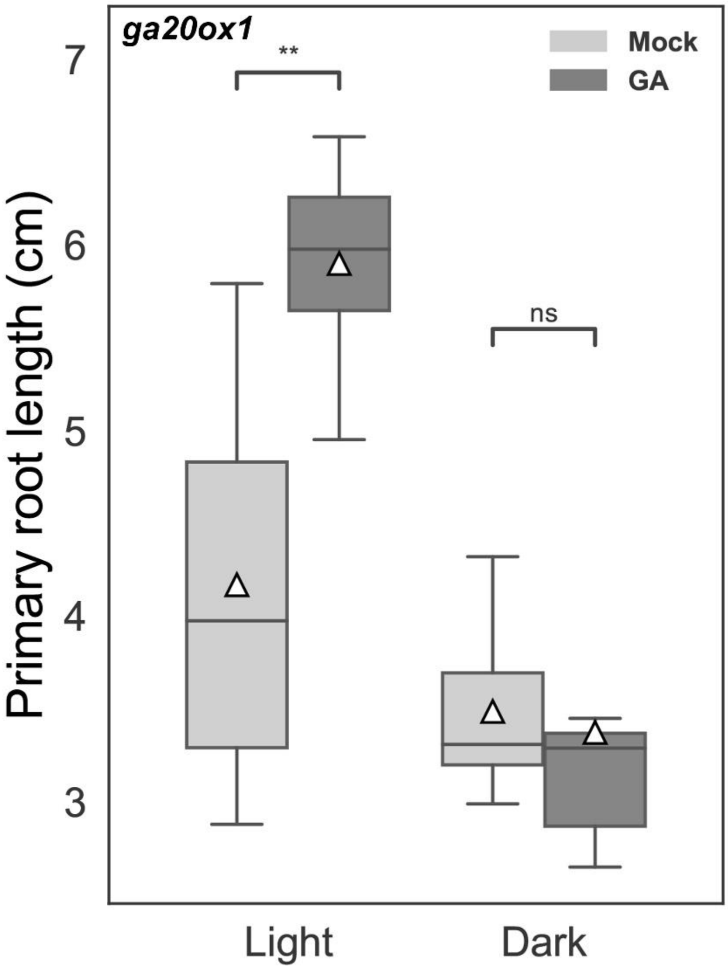


**Fig. S9.** Light is required for GA-induced primary-root elongation in *ga20ox1*. *Ga20ox1* seedlings were grown in agar plates with or without 10µM $GA_3$ and the plates were placed vertically in the light or under dark conditions and after 5 days primary-root length was measured. Values are mean of 16 seedlings ±SE. Stars represent significant differences between respective treatments by Student's t test (P<0.01). ns- not significant.

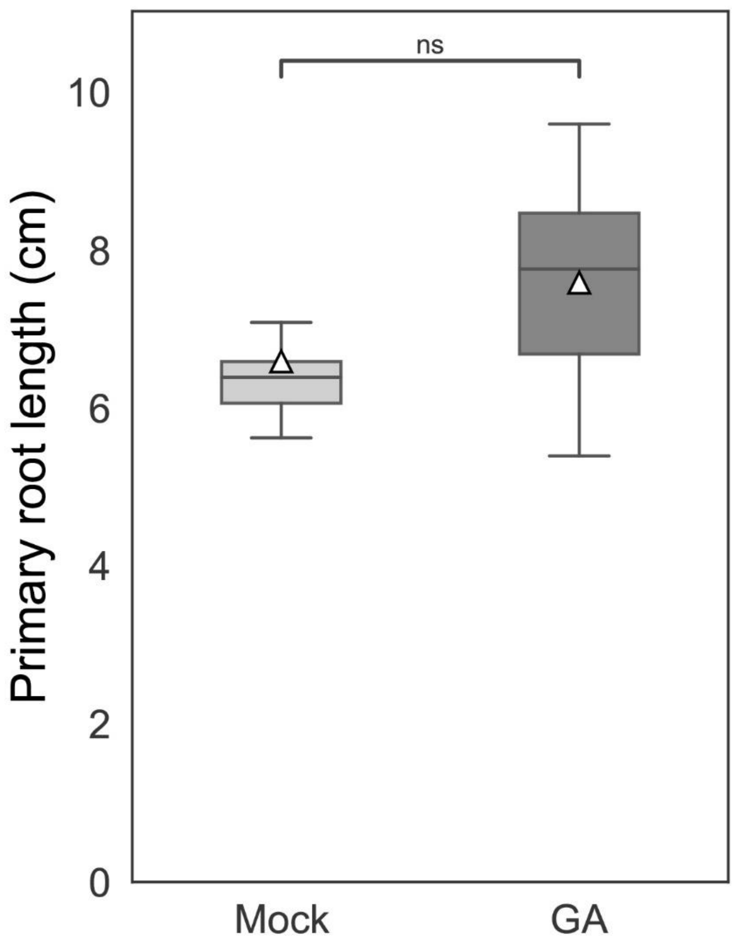


**Fig. S10.** GA has no effect on primary root elongation in M82 seedlings grown in agar plates. M82 seedlings were grown in agar plates for 5 days and then primary-root length was measured. White triangles show the average values (n=10 ±SE). ns- not significant by Student’s t test.

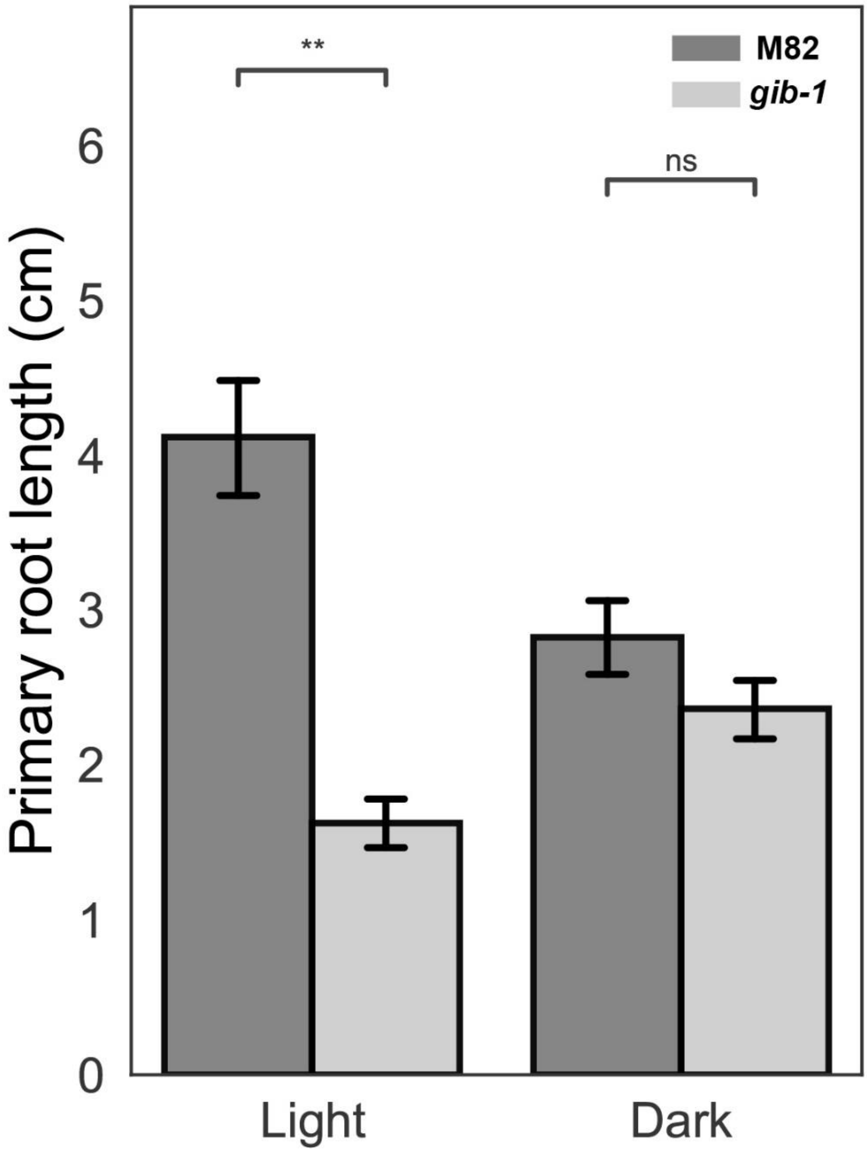


**Fig. S11.** Primary-root length of WT M82 and *gib-1* grown in agar plates under light or dark conditions. WT M82 and *gib-1* seedlings were grown in agar plates. The plates were placed vertically in the light or in dark conditions for 7 days and then the length of the primary roots was measured. Values are mean of 12 seedlings ± SE. Stars represent significant differences between respective treatments by Student's t test (**P<0.01). ns- not significant.

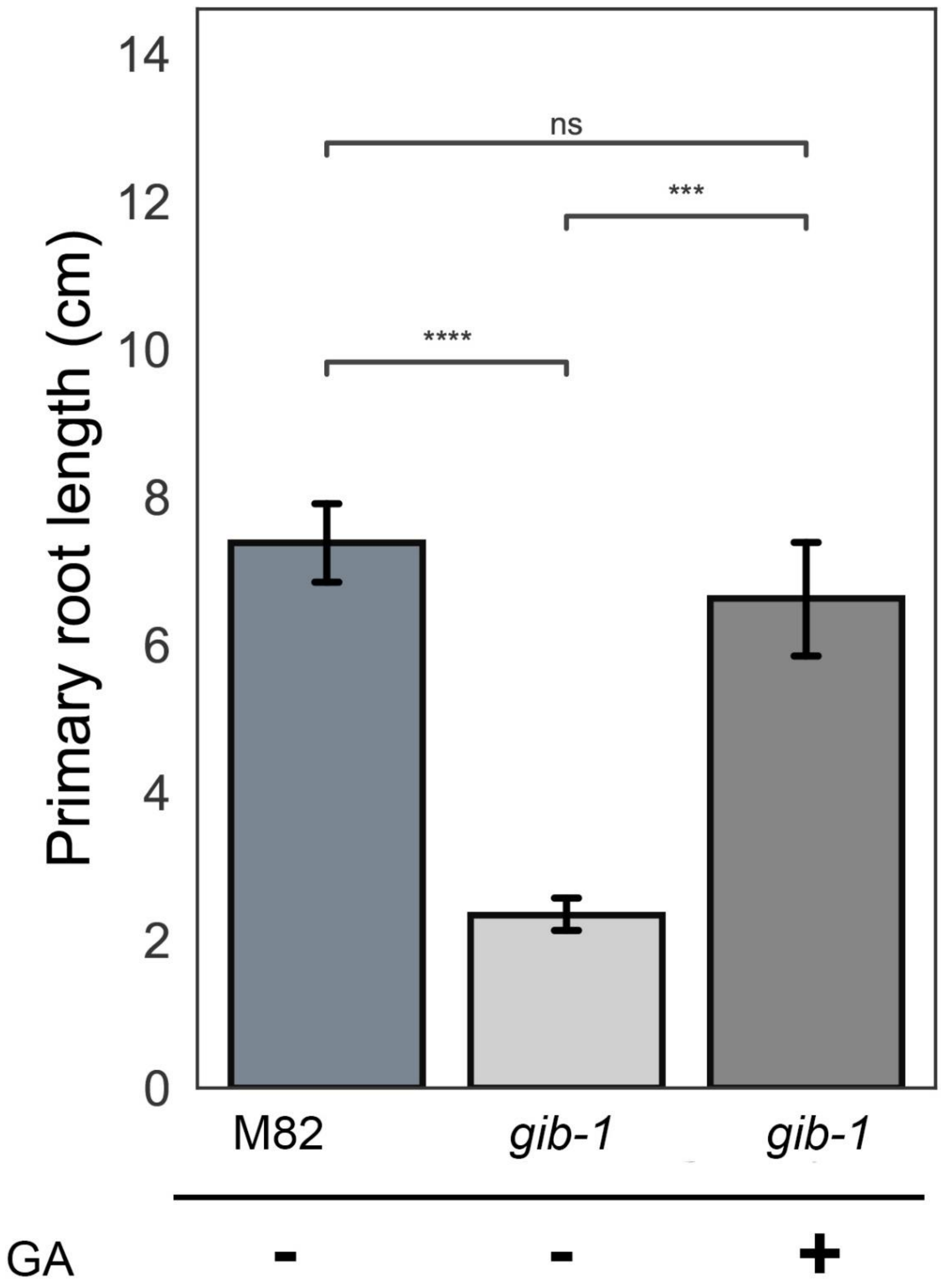


**Fig. S12.** GA application to *gib-1* immediately after germination on vermiculite restored normal primary root elongation. Rescued M82 and *gib-1* embryos were placed on vermiculite and 10μM $GA_3$ was immediately added to *gib-1* and two weeks later the length of the primary roots was measured. Values are mean of 13 seedlings ±SE.

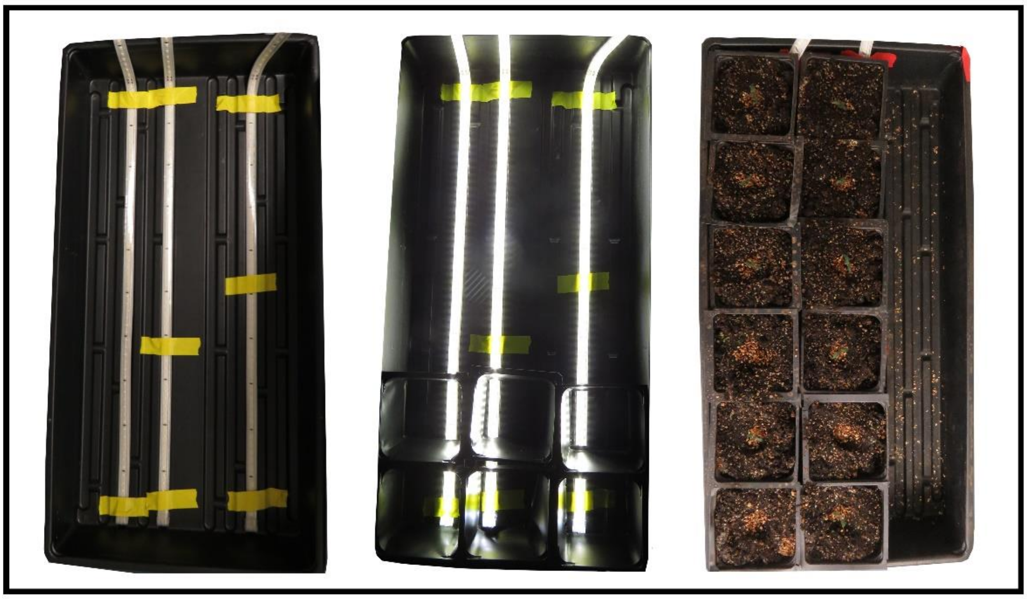

**Fig. S13.** The bottom-lighting growth system. Left- LED lighting strips fixed to the growing tray. Middle- Try with LED lighting strips and pots without their bottom. Right- Try with LED lighting and pots without their bottom, filled with vermiculite and planted *gib-1* seedlings.

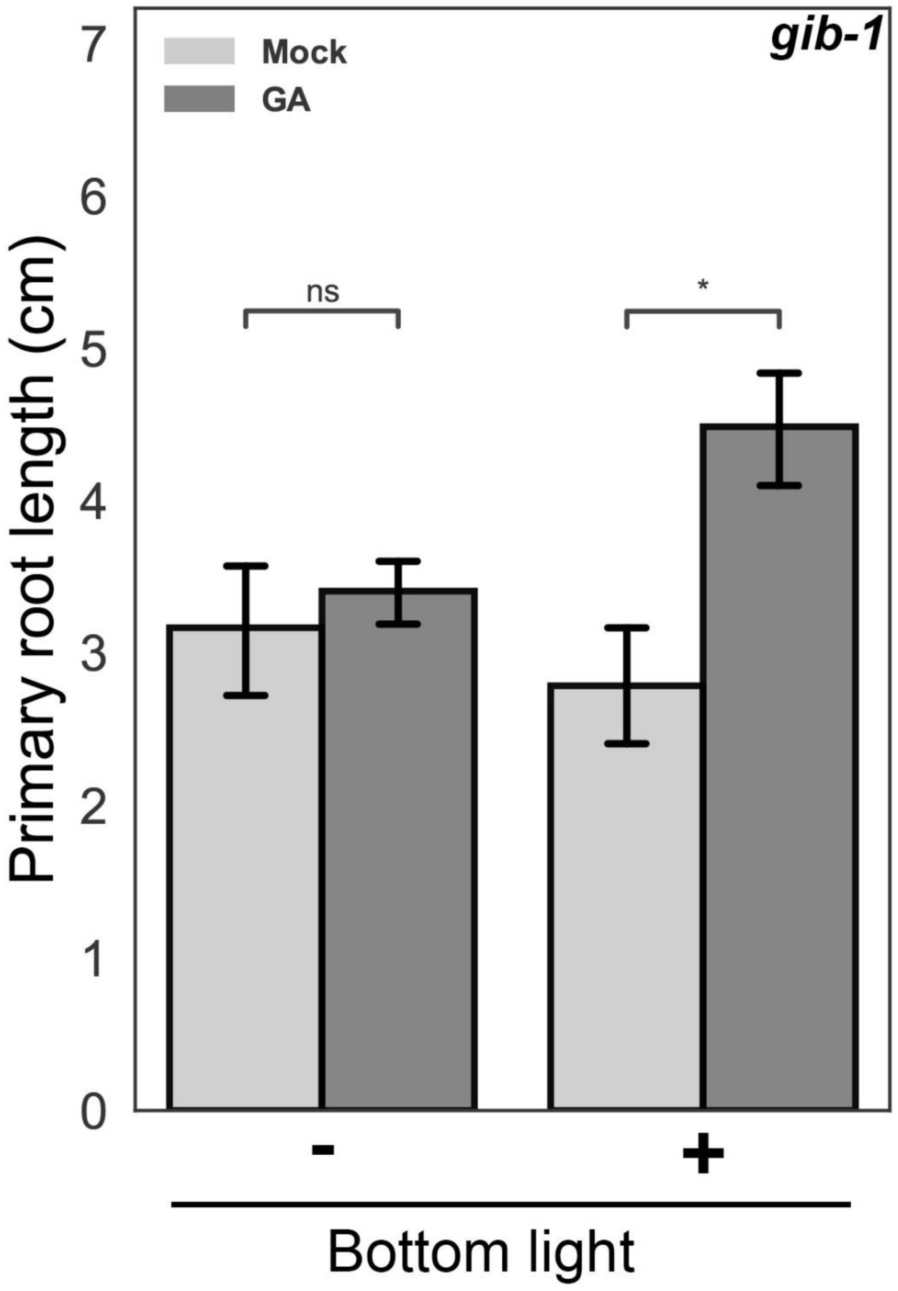


**Fig. S14.** Bottom lightening induced the promoting effect of GA on primary root elongation. *gib-1* embryos were put on the vermiculite in bottomless pots that were placed (or not) on LED lighting strips and two weeks later, when roots were approximately 2.5 cm long, the light was turned on and part of the plants were treated with 10μM $GA_3$. A week later, the length of the primary roots was measured for all treatments. Values are mean of 10 seedlings ±SE. Stars in represent significant differences between respective treatments or lines by Student's t test (*$P < 0.05$). ns- not significant.

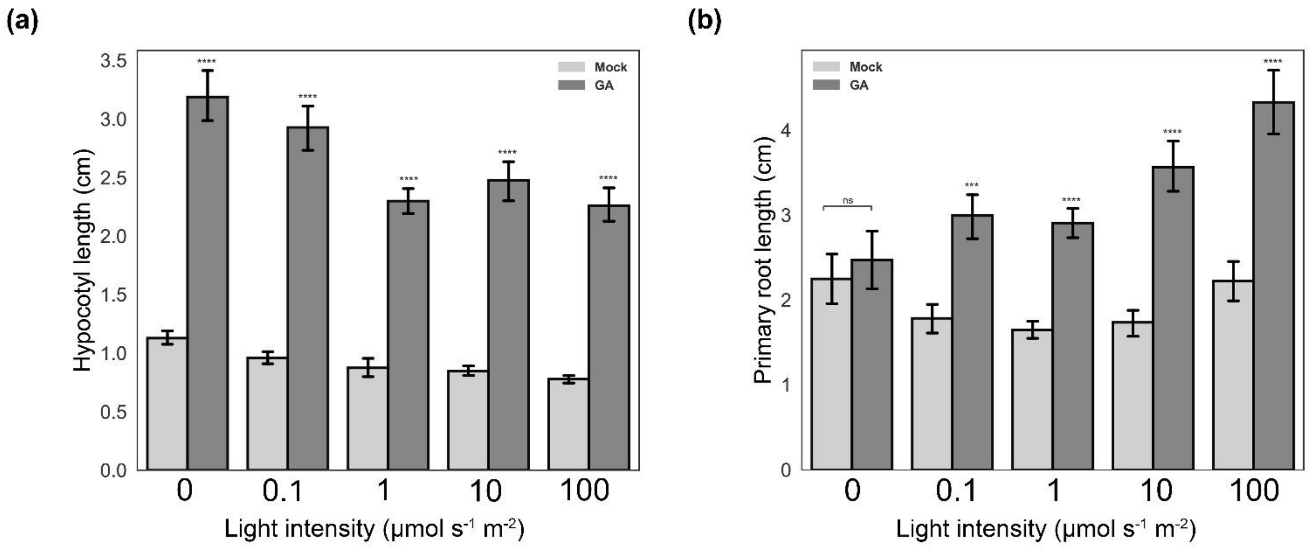


**Fig. S15.** GA-induced hypocotyl and primary root elongation under different light intensity. Rescued *gib-1* embryos were placed on agar plates with or without $GA_3$ in dark or under 0.1, 1, 10 or 100 µmol $m^2$ $s^{-1}$ of white light. After 10 days hypocotyl (a) and primary root (b) length was measured. Values are mean of 9 seedlings ±SE. Stars represent significant differences between respective treatments by Student's t test (***P < 0.001; **P<0.0001). ns- not significant

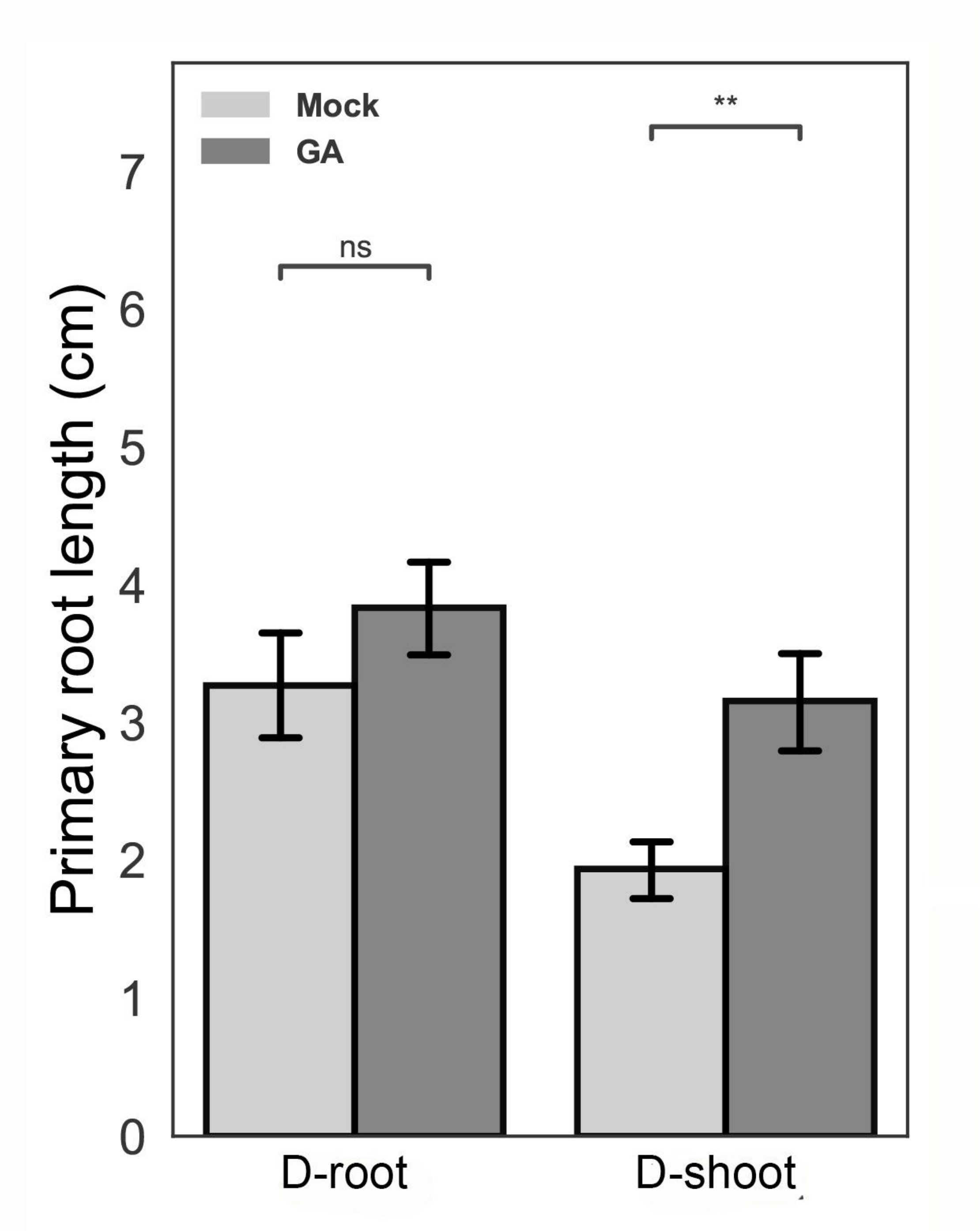


**Fig. S16.** GA promoted root elongation in D-shoot but not in D-root plates. *ga20ox1* seedlings were grown in D-root or D-shoot systems for 7 days and then primary-root length was measured. The average length is presented (n=16 ±SE). Stars represent significant differences between respective treatments by Student's t test (**P<0.01). ns- not significant

**Table S1.** Primers used in this study:

| Gene | Used for | Sequence (5'-3') |
|---|---|---|
| *ACTIN* | RT-qPCR | Forward- GTCCTCTTCCAGCCATCCAT Reverse- ACCACTGAGCACAATGTTACCG |
| *GID1a* | RT-qPCR | Forward- ATGGCAAGAAATAATGAAGCTGT Reverse- CGAAGCTACTTATCTCATCCATG |
| *GID1b1* | RT-qPCR | Forward- GGCTGCTCTTCAATGGGTAA Reverse- TAAAACCTCGACGCCTGATT |
| *ga20ox1* | RT-qPCR | Forward- AGATTGTGTTGGTGGACTTCAA Reverse- TAGCGCCATAAATGTGTCG |
| *ga20ox3* | RT-qPCR | Forward- ACTTTAGGGACAGGGCCTCA Reverse- ACTTGAAGCCCACCAACACT |
